# Systemic stress states are reversed by NPBWR1 inhibition

**DOI:** 10.64898/2026.05.06.722954

**Authors:** Gregor Stein, Philip Müller, Marine Vallet, Emilio Cirri, Lisa Lange, Elisabeth Heller, Julia Winter, Markus H. Gräler, Nico Ueberschaar, Andreas Maurer, Henrik Dobrowolny, Gabriela Meyer-Lotz, Johann Steiner, Olivia Engmann

## Abstract

Chronic stress is widely studied as a brain-centered driver of depression, yet its effects across the body remain unclear. Here, we define chronic stress as a coordinated molecular state across tissues in mice. Using a whole-body proteomic atlas of 13 tissues, we find that stress effects are strongest in peripheral metabolic and endocrine organs, whereas classical stress-associated brain regions show comparatively modest changes. Cross-organ analyses reveal structured, tissue-specific responses rather than uniform shifts. Inhibition of the stress-regulated receptor NPBWR1 reverses both behavioral deficits and proteomic alterations across organs in both sexes, indicating that this state is dynamically modifiable. Integration with human serum proteomics identifies shared, sex-specific signatures. Across tissues, lipid metabolism emerges as a common stress-responsive pathway, confirmed by hepatic lipid remodeling that is normalized by NPBWR1 inhibition. To facilitate exploration of these data, we provide an interactive cross-organ web-based resource. Together, these findings challenge the brain-centric view of stress pathology and define chronic stress as an organism-wide molecular state.

## Introduction

Chronic stress is a major risk factor for psychiatric disorders, including major depressive disorder (MDD) ^1^. It is typically studied as a brain-centered process. However, stress is also linked to cardiovascular, metabolic, and gastrointestinal disease ^2–4^, suggesting effects beyond the central nervous system.

Existing frameworks, including the hypothalamic-pituitary-adrenal (HPA) axis and gut-brain signaling, describe selected inter-organ pathways ^5,6^. Yet these frameworks do not capture how stress reshapes the organism as a whole. Whether stress induces coordinated molecular changes across tissues remains unclear. In particular, it is unknown how stress responses differ between organs, how they vary between sexes, and whether systemic molecular states can be pharmacologically modulated.

Here, we generate a sex-stratified, whole-body proteome atlas across 13 tissues in the chronic variable stress (CVS) mouse model. This approach enables a systematic assessment of tissue-specific and cross-organ stress responses. We find that stress-induced molecular changes are not uniformly distributed, but show the strongest effects in peripheral metabolic and endocrine organs, whereas canonical brain regions show comparatively modest alterations. Cross-organ analyses reveal structured, sex- and tissue-specific coordination of stress responses rather than uniform systemic shifts.

We further test whether this organism-wide state can be modulated pharmacologically. We previously identified the neuropeptide B/W receptor NPBWR1 as a regulator of depression-related phenotypes in the NAc ^7^. Here, we show that systemic inhibition of NPBWR1 using the antagonist CYM50769 reverses both behavioral deficits and molecular alterations across multiple organs in both sexes.

Integration with human serum proteomics reveals shared, sex-specific, reversible signatures between stressed mice and patients with MDD. Across tissues, lipid metabolism emerges as a convergent stress-responsive pathway, supported by hepatic lipidomic remodeling that is normalized by NPBWR1 inhibition.

To facilitate exploration of systemic stress biology, we provide an interactive open-access resource for cross-organ proteomic analysis (https://stress-atlas.com). Together, these findings establish chronic stress as a coordinated organism-wide molecular state that can be pharmacologically reversed.

## Methods

Further information can be found in the *extended methods*.

### Animals

All animal experiments were conducted in accordance with institutional and national guidelines. Experiments performed in Germany were approved by the Thüringer Landesamt für Verbraucherschutz (TLV; license FSU-24-005) and complied with EU Directive 2010/63/EU and the German GenTAufzV. Experiments performed in the United States were approved by the Institutional Animal Care and Use Committee (IACUC; protocol 805959). C57BL/6J mice were bred in-house at the Friedrich Schiller University Jena animal facility or obtained from Janvier Labs (France). Mice were housed in groups of 2 to 5 under a 12 h light–dark cycle and were at least 8 weeks old at the start of experiments. Both sexes were used as indicated.

### Pharmacological treatment

The NPBWR1 antagonist CYM50769 (Tocris, #4948) was dissolved in 1% DMSO in phosphate-buffered saline (PBS), filtered, and administered by intraperitoneal injection (10 ml kg^−1^). Mice received daily injections at doses of 0.4, 4.2, or 41.7 µg kg^−1^. Dose selection was guided by reported in vitro IC_50_ values for NPBWR1 antagonism (120 nM). Control animals received vehicle (1% DMSO in PBS). Injections were performed during the late active phase (−3 to 0 h relative to light onset) to minimize circadian variability.

### Human serum samples and clinical characterization

Serum specimens from acutely ill inpatients with a current episode of major depressive disorder (MDD; n = 82) and matched healthy controls (n = 70) were obtained from the scientific blood biobank of the Department of Psychiatry and Psychotherapy, Otto-von-Guericke-University Magdeburg, Germany. The study was performed in accordance with the Declaration of Helsinki and German laws, and was approved by the local institutional review board of the medical school at Otto-von-Guericke-University Magdeburg (positive ethical vote 110/07 for the “Peripheral biomarkers and T-cell immune response in patients with schizophrenia and mood disorders” project/blood biobanking). Written informed consent was obtained from all participants. Patients were diagnosed with MDD according to DSM-IV-TR criteria and were medication-free at the time of blood sampling (drug-naïve at the first episode, or unmedicated for at least 6 weeks in case of prior antidepressant treatment). Depression severity was assessed using the 21-item Hamilton Rating Scale for Depression (HAMD-21), and psychosocial functioning was assessed using the Global Assessment of Functioning (GAF) scale by trained psychologists within 24 h of hospital admission. Secondary causes of depression were excluded by thorough medical history, physical examination, routine blood analysis (including differential blood cell count, kidney function tests, C-reactive protein, glucose, lipids, liver enzymes, and thyroid hormones), screening for illegal drugs, and cranial magnetic resonance imaging. Exclusion criteria for both patients and controls were a history of immune disease, immunomodulatory treatment, cancer, chronic terminal disease, cardiovascular disorders, diabetes mellitus, substance abuse, and severe trauma. Controls were additionally screened for personal or family history of neuropsychiatric disorders using the Mini-International Neuropsychiatric Interview and excluded in case of disease. Blood was collected at the time of admission (baseline, prior to any in-hospital pharmacotherapy) from fasting participants between 8:00 and 9:00 a.m. into BD Vacutainer™ tubes (Becton Dickinson, Heidelberg, Germany). Serum tubes were allowed to clot for 2 h at room temperature and then centrifuged at 2,000 × g for 10 min. Supernatants were aliquoted into low-protein-binding tubes and stored at −80 °C until analysis. Identical collection and storage procedures were applied to patient and control samples. Demographic and clinical characteristics of the MDD and control cohorts are summarized in Table S2.

### Behavioral tests and CVS

Mice were subjected to a 21-day CVS paradigm consisting of three stressors (tube restraint, tail suspension, or mild foot shocks) applied in a pseudo-randomized sequence without repetition on consecutive days ^8^. Foot shocks (100 per session) were delivered in a shock chamber at randomized intervals. CVS was conducted during the early light phase (0 to 5 h after light onset). Health status was monitored daily and body weight was recorded weekly. The forced swim test was performed as previously described ^7^.

### Conditioned place preference (CPP)

Male mice (10 weeks old) were subjected to a standard two-chamber CPP paradigm with distinct visual and tactile cues ^9^. Baseline preference was assessed one day before conditioning. During conditioning (two sessions per day), mice received cocaine (15 mg kg^−1^, i.p.), CYM50769 (41.7 µg kg^−1^, i.p.), or vehicle in the morning paired with confinement to the less-preferred chamber, and saline (0.9%, 1 ml kg^−1^) in the afternoon paired with the opposite chamber. On the test day, animals were allowed free exploration and time spent in each chamber was recorded. CPP scores were calculated as the change in time spent in the drug-paired chamber between pre- and post-test.

### Proteomics

Tissue samples were lysed under denaturing conditions, reduced, alkylated, and digested with trypsin using S-Trap microcolumns or miniplates, depending on the sample type. Peptides were separated using an Evosep One LC system equipped with a C_18_ analytical column and a 44 min gradient, and analyzed on an Orbitrap Exploris 480 mass spectrometer operated in data-independent acquisition (DIA) mode. MS1 spectra were acquired at high resolution over a mass range of 350–1650 *m/z*, and DIA scans were performed using multiple variable isolation windows with high-energy collisional dissociation. Raw data were processed in Spectronaut v19.9 using the directDIA workflow against the UniProtKB/Swiss-Prot Mus musculus database, controlling the false discovery rate at 1% at both precursor and protein levels. Label-free quantification was performed at the precursor level with local normalization. Differential protein abundance was assessed in R using linear models with Benjamini–Hochberg correction for multiple testing. Proteins with log_2_ fold change > 0.58, and adjusted q < 0.05 were considered significant. Keratin contaminants and unassigned entries were excluded from further analysis.

### Lipidomics

Lipids were extracted from liver tissue using a biphasic methanol/chloroform protocol including internal standards. Following phase separation, the organic phase was collected, dried under nitrogen, and reconstituted for LC–MS/MS analysis. Lipid species were separated on a reverse-phase C_18_ column maintained at 50 °C using a methanol–water gradient containing formic acid, and analyzed on a triple-quadrupole mass spectrometer operated in positive ion mode ^10^. Quantification was performed using Analyst software (v1.6.2) based on internal standards and external calibration curves. Lipid abundances were normalized to tissue input. Statistical analyses were performed using Benjamini–Hochberg correction for multiple testing.

## Results

### Chronic stress predominantly affects the proteome of peripheral organs

To obtain an unbiased, systemic view of stress responses, we performed quantitative proteomics across 13 organs in male and female mice (**Fig. 1a**). We employed a CVS paradigm, which enables sustained stress exposure while permitting sex-stratified analyses, a necessity given the pronounced sex differences in depression ^8^. Across tissues, protein detection ranged from more than 7,700 proteins in highly cellular organs, such as the pituitary gland and testes, to fewer than 700 proteins in serum, reflecting tissue-specific complexity (**Table S1**). Despite comparable sample sizes, stress responses were not uniformly distributed across tissues. Instead, they showed a bias toward peripheral organs.

**Fig. 1.**
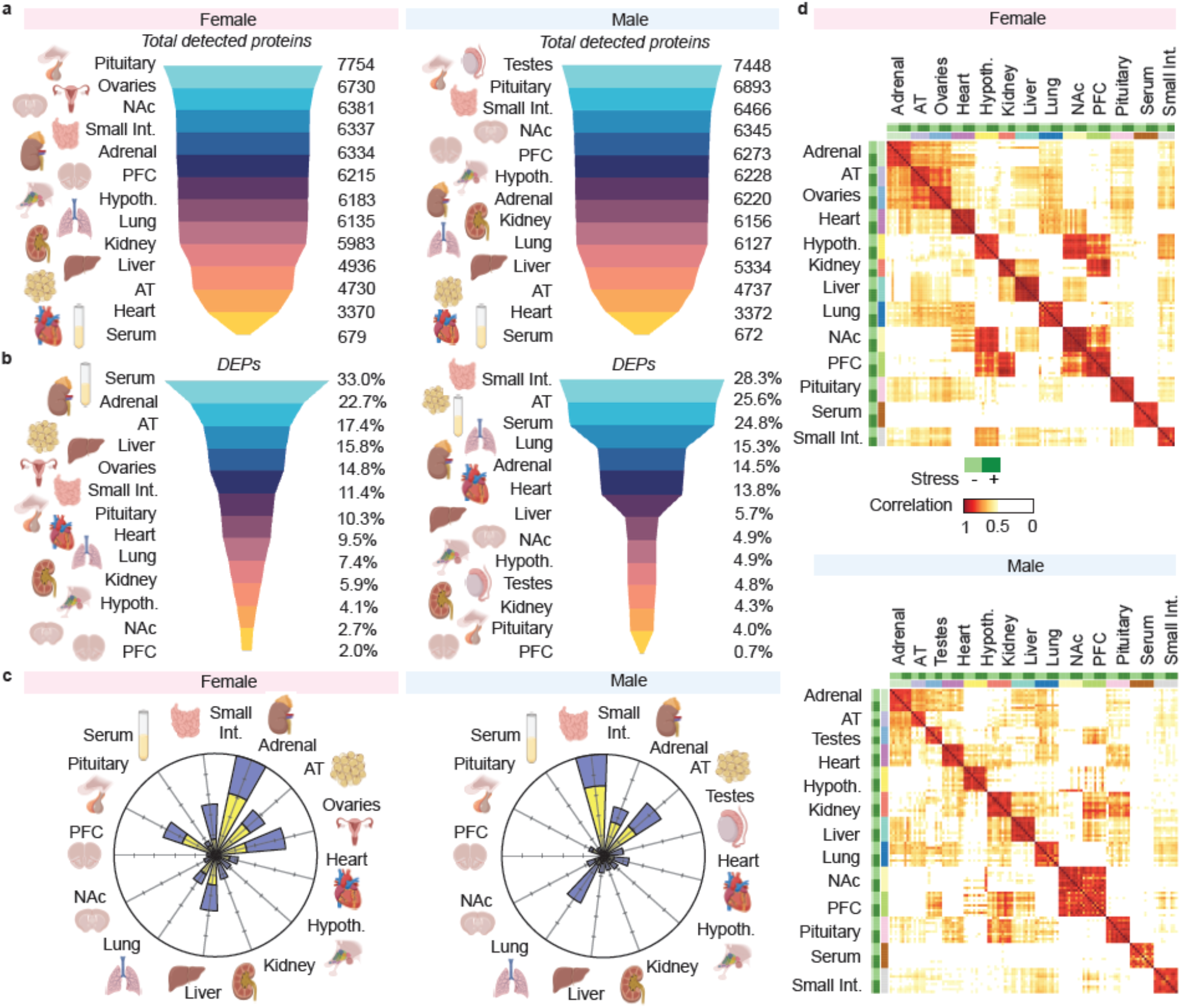
Metrics on organ-specific DEPs in both sexes. **a**, Total quantified DEPs in both sexes differ slightly across tissues, reflecting tissue composition. **b**, DEPs with most stress-induced changes vary drastically across tissues, with the highest percentages in the small intestine and adipose tissue (AT) and smaller percentages in brain regions (NAc, hypothalamus, PFC, pituitary gland). **c**, Petal plots show % of significantly upregulated (blue) vs. downregulated (yellow) DEPs after stress. **d**, Pearson correlations depict the correlation of stress-induced fold changes across tissues. **a-d**) Values represent averages across groups.

In females, the highest proportion of stress-regulated differentially expressed proteins (DEPs) was observed in serum (33.0%), adrenal gland (22.7%), and adipose tissue (17.4%). In contrast, canonical stress-associated brain regions, including hypothalamus, nucleus accumbens (NAc), and prefrontal cortex (PFC), showed the lowest fractions of DEPs (**Fig. 1b**). A similar pattern was observed in males, with the strongest changes in small intestine (28.3%), adipose tissue (25.3%), and serum (24.8%), while NAc, hypothalamus, pituitary gland, and PFC displayed fewer than 5% regulated DEPs.

The direction of protein regulation was highly tissue- and sex-specific (**Fig. 1c**; **Table S1**). In females, the proportion of upregulated proteins ranged from 82.4% in the hypothalamus to 29.3% in adipose tissue. In males, stress responses ranged from predominant upregulation in the lung (80.1%) to a bias toward downregulation in the pituitary gland (20.6% upregulated). These data indicate that stress does not induce a uniform systemic shift, but rather engages distinct, organ-specific regulatory programs.

To assess coordination across tissues, we performed pairwise Pearson correlation analyses of stress-induced proteomic changes (**Fig. 1d**). In females, brain regions (NAc, PFC, and hypothalamus) showed strong concordance, whereas peripheral metabolic organs formed a distinct interconnected cluster. In males, correlations between NAc and PFC were maintained, while associations with the hypothalamus were reduced. In both sexes, the hypothalamus, pituitary gland, and adrenal gland showed moderate correlations with multiple tissues. Serum exhibited limited concordance with other organs.

Together, these data show that chronic stress induces structured proteomic remodeling across the organism. The largest changes occur in peripheral metabolic and endocrine tissues. These responses are highly sex-specific in magnitude, direction, and inter-organ coordination.

### Central and peripheral reversal of stress effects by NPBWR1 inhibition

Having defined stress as an organism-wide molecular state, we next asked whether this state can be pharmacologically reversed. We previously demonstrated antidepressant-like effects of the NPBWR1 inhibitor CYM50769 following intra-NAc administration ^7^. Here, we evaluated whether chronic intraperitoneal delivery of CYM50769 is sufficient to reverse behavioral and organ-wide molecular stress signatures.

Prior to in vivo studies, we assessed compound stability and safety parameters. CYM50769 showed minimal inhibition of hERG channel currents, indicating low predicted cardiac liability. In vitro stability assays demonstrated sustained compound integrity for up to 8 h under physiological conditions, and liposomal clearance rates were slower than reference compounds, suggesting adequate first-pass stability. Positron emission tomography (PET) tracing confirmed rapid brain penetration following systemic administration (**Fig. 2a, b**; **Fig. S1**).

**Fig. 2.**
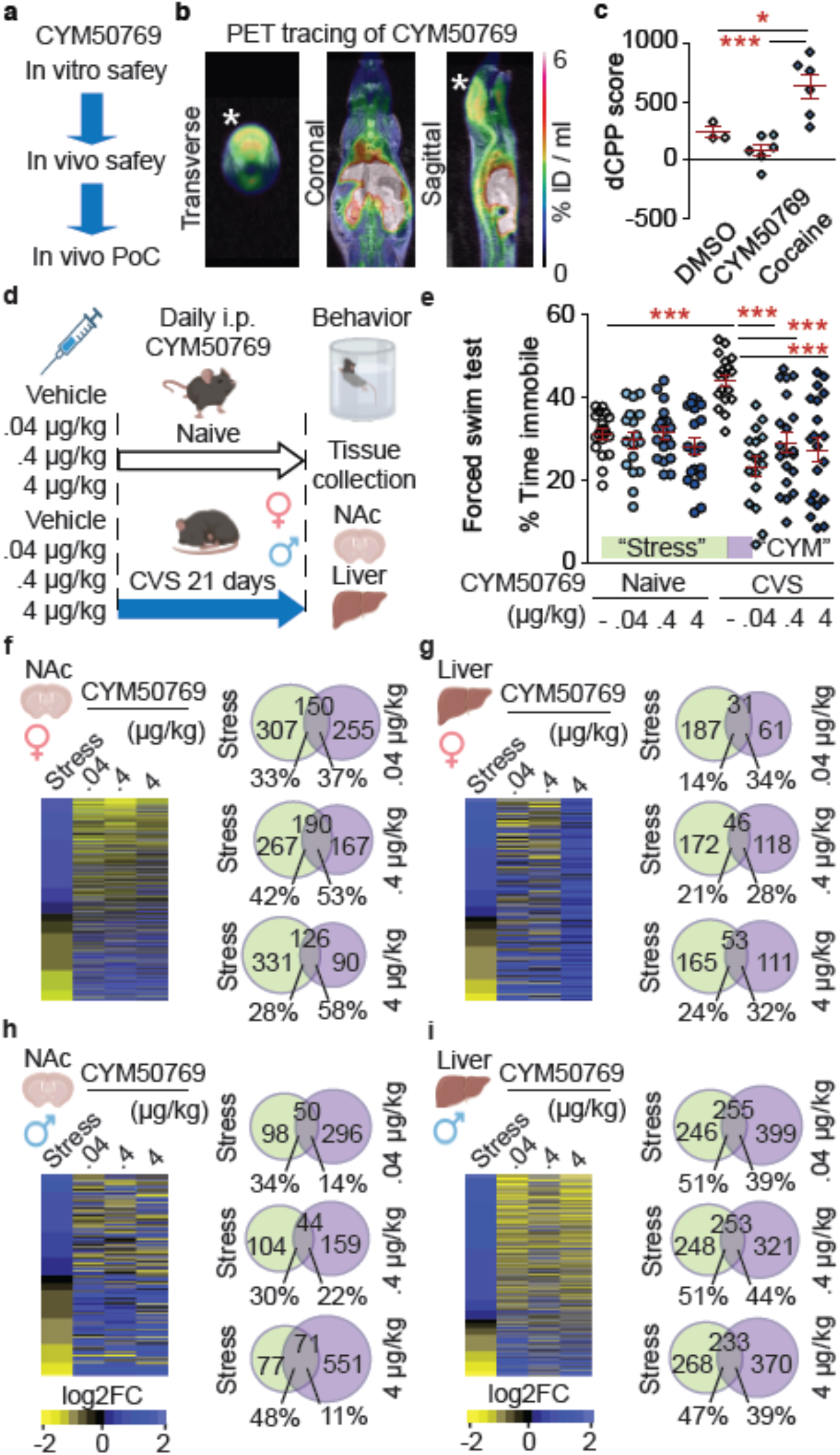
CYM50769 is an efficient and well-tolerated antidepressant-like compound. **a**, Experimental design. **b**, PET tracing shows body-wide distribution of radioactively labelled CYM50769 (asterisk: brain). **c**, Conditioned place preference (CPP) shows that CYM50769 does not induce drug-seeking behavior. n = 3, 6, 6 per group. 1-way ANOVA: F_2,14_ = 13.42, P < 0.001; Tukey *post hoc* test: DMSO vs. CYM50769: n.s.; DMSO vs. cocaine: *P < 0.05; CYM50769 vs. cocaine: ***P < 0.001. **d**, Experimental overview in vivo pilot. **e**, Forced swim test (both sexes pooled) shows that CYM50769 rescues stress-induced increase in immobility time at all doses. n = 19, 17, 19, 19, 18, 17, 20, 20 per group. 2-way ANOVA: drug effect: F_(3,141)_ = 11.56, P < 0.0001; interaction stress x drug: F_(3,141)_ = 8.32, P < 0.0001; Bonferroni *post hoc* test: all doses CYM50769 vs. vehicle within CVS: ***P < 0.001; stress effect within DMSO: ***P < 0.001; all other comp.: n.s. **f-i**, Sex-specific proteomics on organs from mice that were i.p.-injected with 3 doses of CYM50769 versus vehicle under stress / naïve conditions. Heatmaps show the clearest reversal of stress effects by CYM50769 in the lowest dose. Venn diagrams show the largest overlap between stress and CYM50769 in all 3 doses in both sexes. **f, g**, Females. **g, i**, Males. **f, h**, NAc. **g, l**, Liver. “Stress”: DMSO Naïve vs. DMSO CVS; “CYM50769”: DMSO CVS vs. CYM50769 CVS.

Given the role of NPBWR1 in reward-related circuits, we assessed potential intrinsic reinforcing effects using conditioned place preference (CPP). Whereas cocaine induced robust place preference, CYM50769 did not alter reward-seeking behavior (**Fig. 2c**; **Fig. S1**), indicating the absence of intrinsic reward-related side effects.

Mice were then subjected to CVS and treated daily with CYM50769 across a 100-fold dose range (**Fig. 2d**). In both sexes, CYM50769 reversed the stress-induced increase in immobility time in the forced swim test at all doses tested (**Fig. 2e**). Body weight and locomotor activity were unaffected (**Fig. S1**), indicating that behavioral rescue was not driven by nonspecific effects.

To determine whether molecular changes accompanied behavioral recovery, we performed proteomic profiling on the NAc and liver within a pilot cohort of both sexes, testing all doses of CYM50769 versus vehicle. CVS induced robust proteomic alterations in both tissues. CYM50769 partially reversed these stress-induced changes in the NAc compared to the stressed vehicle group. Liver proteomes were also affected by both stress and treatment, albeit with distinct patterns, indicating that peripheral tissues are responsive to NPBWR1 modulation. Across both sexes, the two lower doses produced a reversal in NAc and liver. The highest dose showed less specific effects, suggesting dose-dependent divergence of selective target engagement (**Fig. 2f-i; Fig S1**) ^11^.

Together, these data demonstrate that systemic NPBWR1 inhibition reverses stress-induced behavioral phenotypes and partially normalizes proteomic alterations in both brain and peripheral organs. These findings indicate that the organism-wide stress state is dynamically modifiable by targeted pharmacological intervention.

### Rescue of stress-induced within-tissue proteome changes by NPBWR1 inhibition

We next asked whether systemic NPBWR1 inhibition reverses stress-induced proteomic alterations within individual tissues of both sexes (**Fig. 3**; **Fig. S2, S3**). Across organs, up to 63% (serum) of stress-regulated DEPs overlapped with DEPs altered by CYM50769 treatment (**Fig. 3a; Fig. S3a**). To assess directionality, we examined shared DEPs using fold-change heatmaps. In most tissues and in both sexes, CYM50769 shifted stress-induced protein alterations toward baseline levels (**Fig. 3b, Fig. S3b**), indicating broad within-tissue reversal.

**Fig. 3.**
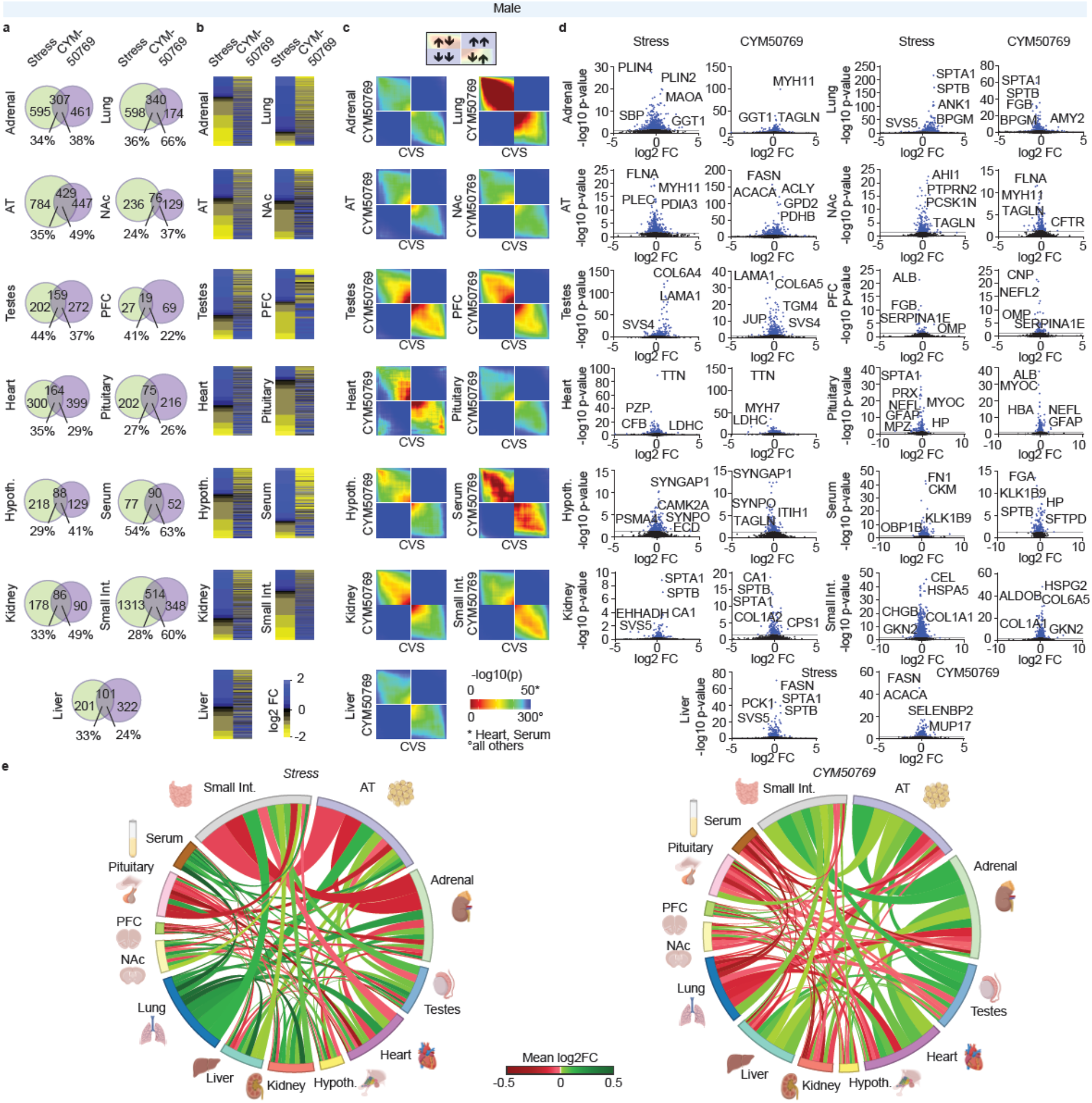
CYM50769 reverses systemic effects of chronic stress in male mice. **a**, Overlap between DEPs significantly changed by stress and CYM50769 (Venn diagrams), shows partial reversal of stress effects by CYM50769. **b**, Heatmaps depict DEPs that are significantly altered by stress and were sorted by log_2_ fold change within the stress group, with the same DEPs depicted for CYM50769 in the right lanes regardless of the adjusted P-value in that group. The heatmaps show an inverse log_2_ fold change within the CYM50769-group across sexes and tissues. **c**, RRHO analysis shows inverse correlation between CVS and CYM50769 across sexes and tissues. **d**, Volcano plots show the distribution of significance and log_2_ fold change altered DEPs. **e**, Chord diagrams reveal stress-induced changes in organ connectomics. These are at least partially reversed by CYM50769. “Stress”: DMSO Naïve vs. DMSO CVS; “CYM50769”: DMSO CVS vs. CYM50769 CVS. AT – adipose tissue.

To quantify global concordance independent of significance thresholds, we performed rank-rank hypergeometric overlap (RRHO) analysis. Across tissues, RRHO plots revealed a consistent inverse association between stress and CYM50769 signatures, indicating that NPBWR1 inhibition broadly counteracts stress-induced proteomic changes (**Fig. 3c, Fig. S3c**). Concordance in the same direction was minimal, supporting a selective reversal of stress signatures rather than non-specific proteome remodeling.

Volcano plots highlighted representative stress-sensitive DEPs that were normalized by CYM50769 treatment. For example, stress-induced upregulation of lipid metabolic regulators in the liver, including FASN, was attenuated following treatment. Similar reversal patterns were observed across multiple organs with sex-specific differences (**Fig. 3d**; **Fig. S2, S3d**). Despite this overall convergence, the underlying molecular responses were highly sex-specific. In several tissues, including the hypothalamus, lung, and serum, stress-induced changes and their reversal showed partially opposing directionality between sexes (**Fig. S4**). These findings underscore the necessity of sex-specific analyses when assessing systemic stress responses. Together, these data show that systemic NPBWR1 inhibition partially restores stress-altered proteomes across both brain and peripheral tissues in males and females.

### Organ-wide reorganization of the stress proteome

Having established that chronic stress alters proteomes across all examined tissues, we next asked whether it also reshapes coordination between tissues and whether such inter-organ relationships are restored by NPBWR1 inhibition. To capture multi-tissue regulation without thresholding, we generated chord diagrams integrating fold-change magnitude and directionality across organs.

In females, stress induced a dense network of inter-organ associations, including expected endocrine links such as pituitary-adrenal connectivity. In addition, coordinated changes emerged between adipose tissue and the adrenal gland, lung and the ovary, and PFC and small intestine, indicating widespread systemic coupling (**Fig. S3e**). Serum and lung displayed an overall downregulation of DEPs shared with other organs, whereas pairs such as kidney-intestine and adrenal-intestine showed a shift towards upregulation. These patterns suggest structured, rather than random, network reorganization.

Treatment with CYM50769 reversed many of these stress-induced associations. Prominent stress-associated connections, including adrenal-small intestine and kidney-small intestine interactions, were attenuated or inverted following treatment (**Fig. S3e**). Serum-associated interactions were particularly sensitive to pharmacological modulation, consistent with its strong stress responsivity.

In males, stress-induced coordination patterns differed markedly. Prominent connections involved adipose tissue-small intestine, adipose tissue-adrenal gland, hypothalamus-adrenal gland, testes-adrenal gland, and small intestine-PFC/NAc links (**Fig. 4e**). In contrast to females, stress-induced coordination in males showed an overall shift towards upregulation across shared DEPs, including interactions involving serum and lung.

**Fig. 4.**
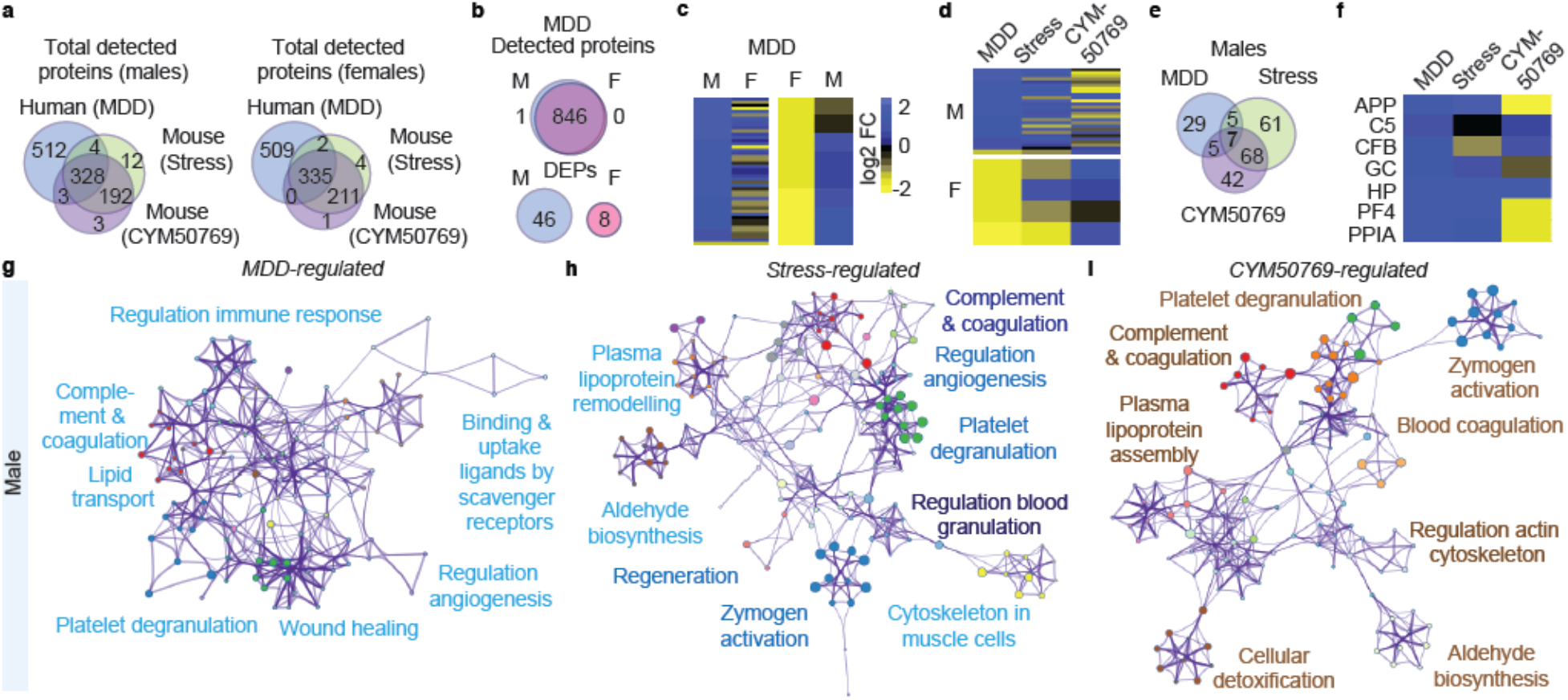
Overlap in serum from MDD patients and mice reveals shared biomarkers. **a**, Venn diagrams reveal that 40% of detected proteins were shared between mice and humans of both sexes. **b**, While detected proteins were nearly identical across genders, major depressive disorder (MDD) was associated with highly sex-specific serum proteome changes, which revealed no overlap between male and female patients. Men showed a >5x higher number of MDD-associated DEPs than women. Therefore, subsequent analyses were conducted separately for both sexes. **c**, In agreement with **b**, no association in directionality of MDD-associated protein changes was detected across sexes. Additionally, the direction of changes differed across genders, with male patients showing predominant upregulation of DEPs (blue) and females showing downregulation (yellow). **e**, Significantly changed DEPs in male patients, male mice that had undergone CVS (DMSO Naïve vs. DMSO CVS) and CYM50769 treatment (CYM50769 CVS vs. DMSO CVS). 7 DEPs were significantly altered in all 3 comparisons. **f**, Heatmap of 7 shared DEPs from **e**: APP, GC, PF4, and PPIA are upregulated during MDD and CVS and downregulated by CYM50769 treatment. Scale as in **c. g-i**, Metascape pathway analysis on male cohorts. Blue: upregulation, brown: downregulation. **g**, Pathways altered in MDD patients are predominantly upregulated and are linked to coagulation, angiogenesis, lipid transport and immune response. **h**, CVS-associated pathways in mice are also predominantly upregulated and, as in patients, relate to blood coagulation, lipoproteins, aldehyde synthesis, and angiogenesis. **i**, CYM50769-regulated pathways are predominantly downregulated and include coagulation, lipoproteins, and aldehyde synthesis.

CYM50769 treatment reshaped these networks and reversed key stress-associated connections (**Fig. 4e**). Together, these analyses show that chronic stress reorganizes inter-organ proteomic coordination in a sex-specific manner and that systemic NPBWR1 inhibition partially restores this organism-wide network structure.

### Sex-specific overlap between stress and depression biomarkers in patient serum

To assess the translational relevance of our findings, we performed proteomics on serum samples from patients with MDD and matched controls (82 patients, 70 controls, **Table S2**). Cohorts were matched for age and body mass index, although male patients exhibited higher smoking rates, and both sexes showed increased waist-to-hip ratios compared to controls. Patients were not receiving antidepressant treatment at the time of sampling.

In total, 847 proteins were detected in human serum, of which 40% overlapped with those found in murine datasets (**Fig. 4a**). Compared to mice, where 28–33% of DEPs were stress-regulated, only a small fraction of proteins was altered in MDD (5% in males, 1% in females), consistent with increased biological variability in human cohorts. Despite this, MDD-associated proteomic changes showed pronounced sex specificity. DEPs exhibited no overlap between male and female patients and differed in directionality, with a bias toward upregulation in males and downregulation in females (**Fig. 4b, c; Fig. S5**). Despite the substantially higher variability in human samples, this pattern mirrored sex-specific stress responses observed in mice (**Fig. 4d**). 17 DEPs were shared between male MDD patients and stressed male mice (37% of MDD-associated DEPs), of which seven were also significantly altered by CYM50769 treatment (**Fig. 4e**). Four of these DEPs showed stress- or disease-associated upregulation that was reversed by CYM50769 in mice. Among these, amyloid precursor protein (APP) exhibited the most prominent changes (**Fig. 4f**). Additional shared DEPs included platelet factor 4 (PF4), peptidyl-prolyl cis–trans isomerase A (PPIA), and vitamin D-binding protein (GC), which are linked to inflammation, coagulation, and vascular function.

Consistently, metascape pathway analysis revealed an enrichment of platelet degranulation, complement and coagulation cascades, and angiogenesis in both male MDD patients and stressed male mice. These pathways were reversed by CYM50769 in the animal model (**Fig. 4g-i**). Together, these data identify sex-specific serum proteomic signatures that are shared between human depression and experimental stress, and suggest that NPBWR1 inhibition targets conserved systemic pathways linked to inflammation and vascular biology.

### Lipid metabolism emerges as a cross-organ stress response axis

Given the coordinated inter-organ responses induced by stress, we next asked whether common biological pathways are shared between tissues. Enrichment analyses identified lipid metabolism as one of the most consistently altered pathways following chronic stress and CYM50769 treatment (**Fig. 5a, b**), in agreement with pathway signatures observed in serum (**Fig. 4g-i**).

**Fig. 5.**
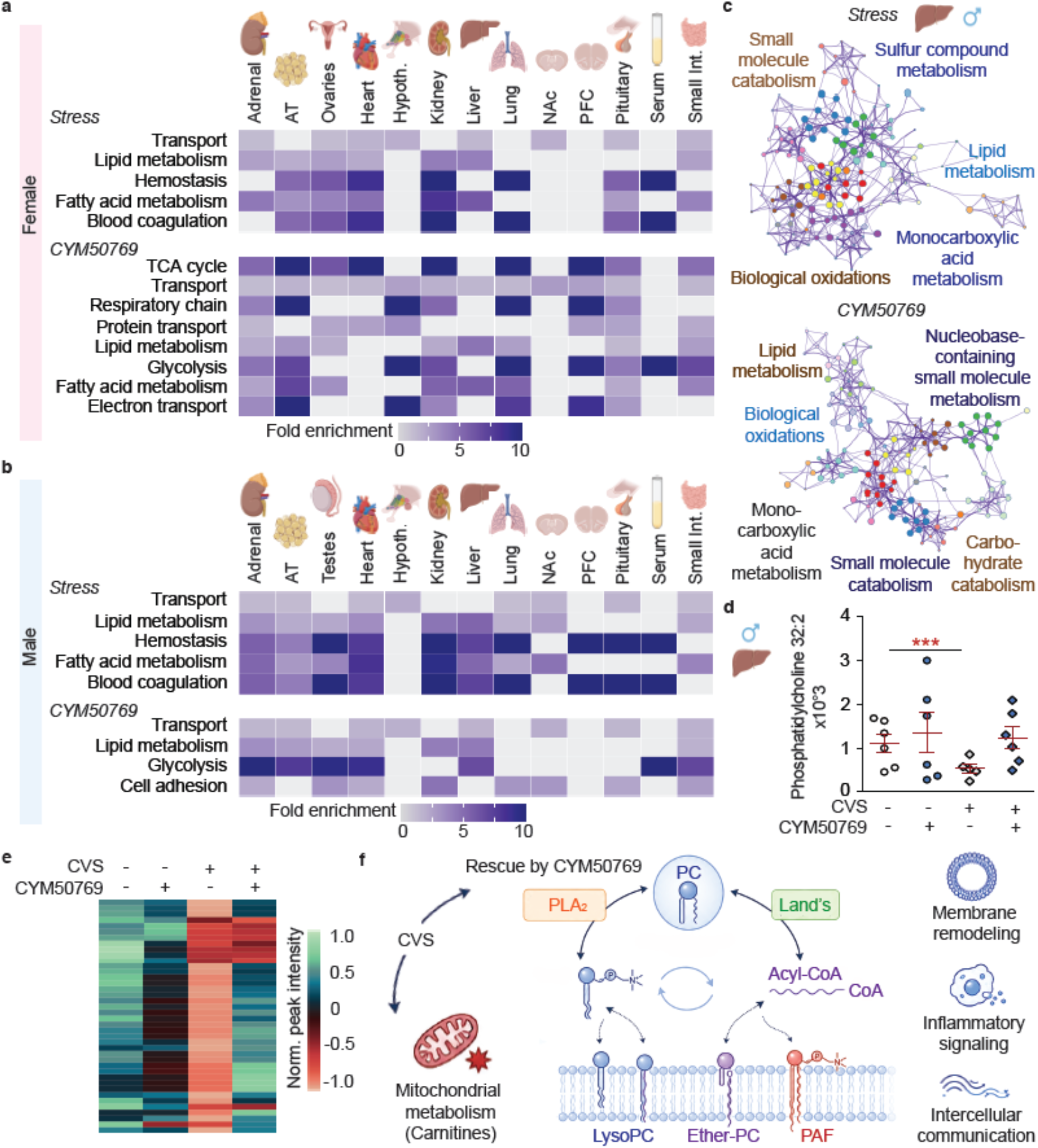
Lipid metabolism is affected by stress and reversed by CYM50769. **a, b**, GO-term analysis of downregulated DEPs after stress in all organs shows a strong implication of lipid metabolism. **a**, Females, **b**, Males. **c**, Metascape analysis in the liver of stress- and CYM50769-regulated DEPs in males. Blue: up, yellow: down. Note the inverse effects of stress and CYM50769 on lipid metabolism, biological oxidations, and small-molecule catabolism. **d**, Lipidomics reveal that phosphatidylcholine 32:2 is reduced in the male liver by stress and rescued by CYM50769. n = 6, 6, 5, 6 per group. 2-way ANOVA: Effect of CYM50769: F_(1,19)_ = 4.57, P < 0.05. Bonferroni *post hoc* test: Stress effect within DMSO: ***P < 0.001; all other comp. n.s. **e, f**, Untargeted metabolomics. Only metabolites with MS/MS data with a full spectral match in the SIRIUS database are shown and retained in the list of 40 curated metabolites. **e**, CVS increases detected polar species, which contain predominantly lipids (lanes 1 vs. 3). CYM50769 in naïve mice induced few changes (lane 2). CYM50769 in the CVS group rescues predominantly phosphatidylcholines but not changes in carnitine signaling (blue cluster in lane 4), see **Fig. S7. f**, CVS predominantly phosphatidylcholine (PC) and carnitine signaling. PC signaling is rescued by CYM50769. Altered PC species suggest remodeling via Land’s cycle (Land’s, reacylation) and Phospholipase A_2_ (PLA_2_). PAF – platelet-activating factor. Individual data points, standard deviation +/−s.e.m., are shown. AT = Adipose tissue.

In females, lipid metabolic processes were broadly affected across endocrine and metabolic tissues, extending beyond classical metabolic organs and indicating organism-wide involvement (**Fig. 5a**). In males, lipid-related pathways, including fatty acid metabolism, lipid metabolism, and hemostasis, were enriched across multiple peripheral organs, particularly liver, adipose tissue, adrenal gland, and small intestine (**Fig. 5b**).

To assess pathway-level coordination, we performed network analyses using Metascape. In male liver, stress-induced networks centered on lipid metabolism, monocarboxylic acid metabolism, and biological oxidation. NPBWR1 inhibition reshaped these networks and reversed lipid-associated pathways (**Fig. 5c**). A similar reorganization was observed using iPath mapping (**Fig. S6**).

To validate these findings at the metabolite level, we performed targeted lipidomics and untargeted metabolomics in liver tissue. Lipidomics showed that phosphatidylcholine (32:2), was decreased by stress and normalized following CYM50769 treatment (**Fig. 6d**). Likewise, metabolomics on liver tissue revealed a stress-induced reduction in lipids matching the increased lipid metabolism in **Fig. 6c**.

**Fig. 6.**
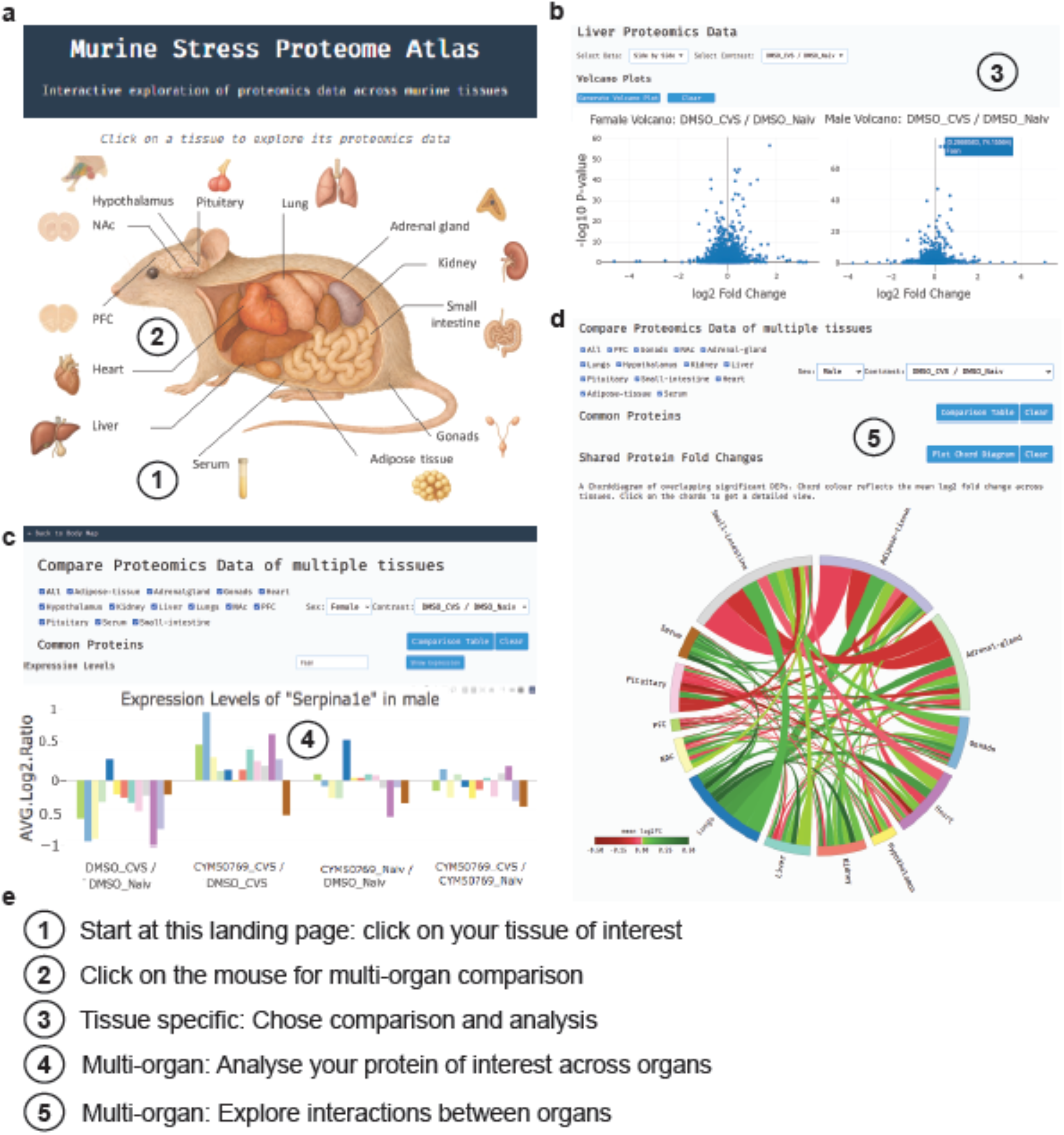
Interactive murine stress proteome atlas for cross-tissue analysis. **a**), Landing page of the atlas on https://stress-atlas.com. Clicking on the mouse will initiate analysis across multiple organs. Alternatively, individual tissues can be selected for proteomic exploration. **b**, The tissue-specific analysis view allows a selection of analyses, including volcano plots, heatmaps, and iPath3 analysis of differential protein abundance between selected experimental conditions (e.g., DMSO Naive versus DMSO CVS). **c**, Multi-organ comparison module enabling visualization of protein abundance changes across tissues. Bar plots show log_2_ fold change ratios for a selected protein across multiple organs and experimental conditions. **d**, Cross-tissue integration of proteomic changes using chord diagrams to illustrate shared and tissue-specific protein regulation. Connections indicate overlapping DEPs between organs, with color coding reflecting the directionality of regulation. **e**, Workflow overview of the platform: (1) selection of tissue of interest from the landing page; (2) initiation of multi-organ comparison via the mouse schematic; (3) tissue-specific differential analysis; (4) cross-tissue visualization of individual protein expression; and (5) exploration of inter-organ relationships.

Of 100 significantly altered features, 40 metabolites were identified by spectral matching analysis with public databases. CYM50769 restored 80% of these curated metabolites within the stressed group, which are predominantly associated phosphatidyl choline signaling (**Fig. 6e, f; Fig. S7**). 8 carnitine-related metabolites were not reversed by CYM50769. Together, these data identify lipid metabolism, particularly phosphatidyl choline signaling, as a conserved cross-organ axis of the stress response and show that systemic NPBWR1 inhibition partially restores this metabolic network.

### An interactive resource for systemic stress proteomics

To enable exploration of organism-wide stress responses, we developed an interactive web-based platform (https://stress-atlas.com). The resource allows tissue-specific interrogation of proteomic changes across experimental conditions, including stress and NPBWR1 inhibition (**Fig. 7a-c**). Users can visualize differential protein expression within individual tissues and compare conditions using interactive volcano plots and heatmaps. In addition, the platform supports multi-organ analyses, enabling the assessment of protein abundance changes across tissues and the identification of organ-restricted versus systemically regulated DEPs.

To facilitate the analysis of inter-organ coordination, the resource includes interactive chord diagrams that capture shared protein regulation across tissues. Specific organ–organ connections can be queried to retrieve underlying protein sets, allowing targeted exploration of cross-organ relationships (**Fig. 7c-e**).

Pathway-level analyses are integrated to enable interrogation of stress- and treatment-associated biological processes, including enrichment-based and pathway-mapping approaches. This allows users to assess how disease-relevant pathways are modulated across organs in response to chronic stress and pharmacological intervention.

Together, this resource provides a framework for systematic exploration of stress-induced proteomic remodeling and its pharmacological modulation across the organism.

## Discussion

Chronic stress is typically viewed as a brain-centered process. Our data challenge this view by showing that stress is encoded as a coordinated molecular state across the organism. Stress-induced proteomic changes were comparatively modest in canonical stress-associated brain regions, whereas peripheral metabolic and endocrine tissues exhibited substantially stronger alterations. This difference may reflect cellular heterogeneity in the brain, where stress-responsive changes are confined to specific neuronal populations and diluted in bulk analyses, whereas peripheral tissues undergo broader metabolic adaptations.

Across mouse tissues and human serum, stress responses were highly sex-specific. Both the identity and directionality of protein changes differed between sexes, consistent with previous observations in human brain tissue ^8^. These findings highlight the importance of incorporating sex as a biological variable in mechanistic studies, biomarker development, and clinical trial design. Despite these differences, NPBWR1 inhibition largely restored stress-induced alterations in both sexes, indicating that sex-divergent molecular states remain pharmacologically tractable. Integration with human serum proteomics identified a subset of shared, sex-specific signatures between experimental stress and MDD. Although only a subset of systemic changes was reflected in serum, likely due to technical and biological variability, key pathways related to inflammation, coagulation, and vascular function were consistently affected. These findings suggest that peripheral biomarkers may capture specific components of the systemic stress response but require validation in larger and clinically stratified cohorts.

A central mechanistic question is how systemic inhibition of NPBWR1 produces widespread proteomic normalization despite its restricted expression pattern. In mice, NPBWR1 is primarily expressed in brain regions such as the NAc, PFC, hypothalamus, and pituitary gland. Therefore, the systemic rescue observed here, likely reflects secondary network-level effects originating from central circuits rather than direct receptor engagement in peripheral tissues. Given the integrative role of the NAc in stress and neuroendocrine signaling, modulation of this node is likely to propagate through hormonal and autonomic pathways, reshaping peripheral proteomic states.

These findings suggest that systemic molecular changes following antidepressant intervention are unlikely to represent off-target effects, but rather components of the therapeutic mechanism. This perspective extends current models of antidepressant action from a primarily neural framework to a systems-level view of coordinated organism-wide regulation.

Lipid metabolism emerged as a conserved pathway across tissues and sexes, supported by proteomic and lipidomic analyses. This is consistent with growing evidence linking metabolic dysregulation to stress-related disorders. Together with alterations in coagulation and platelet activation, these findings suggest that chronic stress engages pathways associated with cardiometabolic risk, potentially contributing to common comorbidities such as cardiovascular disease and fatty liver.

Several limitations should be considered. First, the study is largely based on mouse models, and although integration with human serum supports translational relevance, broader cross-species validation is required. Second, bulk tissue proteomics does not resolve cell-type-specific changes, particularly in the brain. Future studies using spatial or single-cell approaches may refine the resolution of stress-responsive circuits.

In summary, we define chronic stress as a coordinated organism-wide molecular state and show that targeted modulation of a central stress-responsive receptor can partially restore systemic proteomic balance.

## Acknowledgments

This project was made possible by funding from the Carl-Zeiss-Foundation (IMPULS #P2019-01-0006, O.E., G.S., L.L.), from the BMBF Go Bio Initial Grant Sondierungsphase (#16LW0589, O.E.), a LIFE-Connect grant by Friedrich Schiller-University Jena (O.E., N.U.), and NIH R01DA052465 (E.A.H.). M.V. is supported by the Deutsche Forschungsgemeinschaft (DFG, German Research Foundation), SFB1127 ChemBioSys, # 239748522.

We thank Dr. Martina Korfei, Dr. Hussain and Maria Hetzner (all Friedrich Schiller University Jena) for helping with stress induction and i.p.-injections. We furthermore thank Prof. Anna Junker (Werner Siemens Imaging Center, University of Tübingen, Germany) for her contribution to the PET tracing experiment.

## Contributions

G.S., L.L. P.M., and O.E. performed the stress induction, injections, animal scoring, and behavioral tests. O.E., G.S., P.M. and E.C. performed the proteomics analysis. P.M. designed the website. G.S. and L.L. double-checked the data. O.E. planned the experiments and wrote the manuscript. A.M. conducted the synthesis of radioactively labeled CYM50769 and PET tracing. N.U. and G.S. performed stability analyses by HPLC. J.W. and E.A.H. conducted conditioned place preference. M.G. did the lipidomics testing. M.V. and N.U. performed and analysed metabolomics data. J.S. and G.M-L. obtained and characterized human serum samples., H.D. managed the patient database.

