## Supplemental information for "Systemic stress states are reversed by NPBWR1 inhibition"

### Supplementary material

#### Supplementary methods

**hERG channel inhibition assay.** Potential cardiac liability was evaluated by measuring inhibition of the human ether-à-go-go-related gene (hERG) potassium channel using a patch-clamp–based assay (WuXi AppTec). CYM50769 was tested across a range of concentrations, and current inhibition was quantified relative to vehicle control.

**Liver microsomal stability assay.** Metabolic stability of CYM50769 was assessed in vitro using liver-derived microsomal preparations (WuXi AppTec). The compound was incubated at 37 °C in the presence of NADPH-regenerating system. Aliquots were collected at defined time points. Reactions were quenched with organic solvent containing internal standard and analyzed by LC–MS/MS. The percentage of parent compound remaining over time was used to calculate intrinsic clearance and half-life.

**Chemistry.** All reagents and solvents were obtained from commercial suppliers and used without further purification unless otherwise stated. Compound identity and purity (>95%) were confirmed by HPLC, LC–MS, and NMR spectroscopy. Analytical HPLC was performed using a Waters HPLC system equipped with a photodiode array detector. LC–MS measurements were carried out on a Waters Micromass ZQ 4000 mass spectrometer. NMR spectra were recorded on a Bruker Avance III HDX 400 spectrometer and processed using MestReNova software. Precursor 5 for radiosynthesis was synthesized in a multistep procedure starting from 4,5-dichloropyridazin-3(2H)-one involving fluorene coupling, phenoxy substitution, and deprotection reactions (Scheme 1). Final compounds were purified by silica gel chromatography and characterized by HPLC, LC–MS, and NMR spectroscopy.

**Radiochemistry.** [$^{11}$C]CO2 was produced on a PETtrace 890 cyclotron (GE Healthcare, Uppsala, Sweden) and converted to [$^{11}$C]CH3I using a Tracerlab FX MeI module (GE Healthcare) according to the manufacturer's instructions. Radiolabeling, purification, and formulation were performed using a Tracerlab FX M module (GE Healthcare).

**Radiosynthesis of [$^{11}$C]CYM50769.** [$^{11}$C]CH3I was bubbled through a solution containing precursor 5 (1 mg) dissolved in DMF (300 µL) in the presence of 7.5 µL 5 M NaOH at −25 °C. The reaction mixture was heated to 80 °C for 3 min and subsequently cooled to 40 °C. The crude reaction mixture was purified by semipreparative HPLC using a Luna C18(2) column (Phenomenex) with 75% acetonitrile in 0.1 M ammonium acetate at a flow rate of 6 mL min−1. The product-containing fraction was diluted with water, trapped on a Sep-Pak C18 cartridge, eluted with ethanol, and reformulated in PBS.

Quality control was performed by analytical HPLC using a Luna C18(2) column with 80% acetonitrile in 0.1% aqueous trifluoroacetic acid at an isocratic flow rate of 1 mL min−1. The identity of [^11C]CYM50769 was confirmed by co-elution with the nonradioactive reference compound. Radiochemical yield was 332 ± 248 MBq (decay-corrected yield: 2.27 ± 1.73%) with a molar activity of 181.3 ± 43.9 GBq µmol−1 (n = 5).

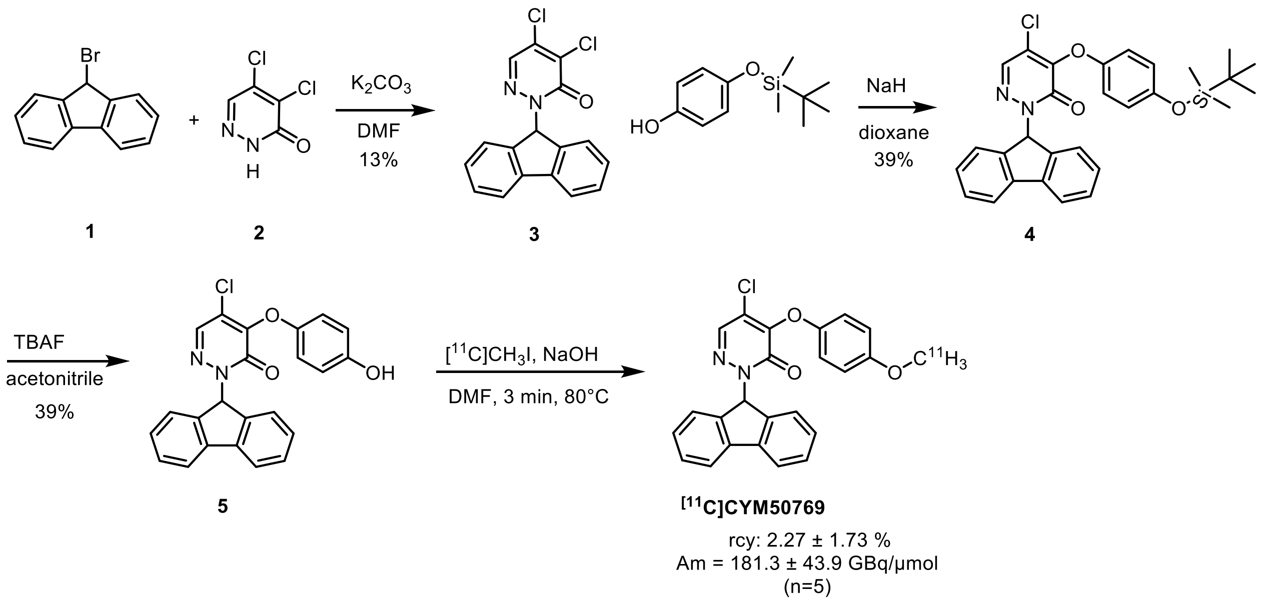


Scheme 1: Synthesis of [$^{11}$C]CYM50769.

**PET/MRI data acquisition and analysis.** Anesthesia was maintained with 1.5% isoflurane in 100% oxygen at a flow rate of 0.8 L min$^{-1}$. Dynamic PET data were acquired over 60 min and reconstructed into 39 time frames (12 × 5 s, 6 × 10 s, 6 × 30 s, 5 × 60 s, and 10 × 300 s) using the OSEM3D reconstruction algorithm. For attenuation correction, a 13 min transmission scan using a cobalt-57 point source was acquired after PET imaging.

Whole-body PET scans were co-registered to anatomical MR images acquired sequentially on a 7T small-animal MRI scanner (Bruker BioSpin GmbH, Ettlingen, Germany). Anatomical MR images were obtained using a TurboRARE T2 sequence (TR/TE 800/35.1 ms; field of view 37.4 × 85.8 × 22.8 mm; matrix size 144 × 256 × 92) with a rat whole-body coil (Bruker BioSpin GmbH). Volumes of interest (VOIs) for blood, muscle, liver, gall bladder, brain, and kidneys were manually defined based on MR anatomical information using PMOD software (v3.2; PMOD Technologies, Zürich, Switzerland). Tissue time–activity curves were calculated and expressed as %ID ml$^{-1}$.

**Proteomics Sample Preparation and LC-MS DIA Analysis.** Mouse serum samples (n = 5) were depleted of abundant serum proteins using the ProteoSpin™ Abundant Serum Protein Depletion Kit (Norgen Biotek) according to the manufacturer's instructions. Proteins were precipitated with ice-cold acetone overnight at −20 °C, washed with 80% acetone, air-dried, and resuspended in digestion buffer (1 M guanidine HCl in 0.1 M HEPES, pH 8). Samples were sonicated and sequentially digested with LysC and trypsin. Digests were acidified with trifluoroacetic acid, desalted using Oasis HLB μElution plates (Waters), dried under vacuum, and stored at −20 °C until LC–MS analysis.

Mouse tissues (excluding serum) were homogenized in PBS containing protease and phosphatase inhibitors (Roche) using a bead homogenizer. Proteins were lysed in SDS-containing buffer, reduced with DTT, alkylated using iodoacetamide, and processed using S-Trap microplates according to the manufacturer's instructions. Following tryptic digestion,

peptides were eluted, dried under vacuum, and reconstituted in MS buffer supplemented with iRT standards. Samples were loaded onto Evotips (Evosep) according to the manufacturer's protocol.
Mouse peptides were separated using an Evosep One system (Evosep, Denmark) coupled to an Orbitrap Exploris 480 mass spectrometer (Thermo Fisher Scientific, Germany). DIA acquisition was performed over a mass range of 350–1650 *m/z* using 40 variable isolation windows. MS1 spectra were acquired at 120,000 resolution and MS2 spectra at 30,000 resolution using higher-energy collisional dissociation.
Human serum samples were depleted of abundant serum proteins as described for mouse serum. Following acetone precipitation, samples were processed using an SDS/S-Trap workflow and analyzed using a Vanquish Neo UHPLC system coupled to an Orbitrap Astral mass spectrometer (Thermo Fisher Scientific). DIA acquisition was performed in positive ion mode using dynamic isolation windows. MS1 spectra were acquired at 240,000 resolution over a mass range of 380–980 *m/z*.
Raw DIA data were analyzed using the directDIA pipeline in Spectronaut v19.9 (Biognosys, Switzerland) using default settings except for local normalization and protein-group-specific filtering. Data were searched against SwissProt mouse or human databases supplemented with common contaminants. Identifications were filtered to a false discovery rate (FDR) of 1% at peptide and protein levels. Protein quantification tables were exported for downstream analysis and visualization in RStudio using custom R scripts. Significant proteins were defined using a $\log_2$ fold-change cutoff of 0.58 and $q < 0.05$ following Benjamini–Hochberg correction. Human serum data were additionally analyzed using MSstats after removal of sparse precursors and filtering for missing values. Keratin contaminants and unassigned protein entries were excluded from downstream analyses.
Raw proteomics datasets were deposited in the MassIVE repository under accession numbers MSV000101519, MSV000101596, MSV000101604, MSV000101599, MSV000101600, MSV000101598, MSV000101595, MSV000101597, MSV000101543, MSV000101601, MSV000101606, MSV000101537, MSV000101631, MSV000101536, MSV000101605, and MSV000101644.

**Lipidomics**. Lipid profiling was performed using liquid chromatography-tandem mass spectrometry (LC–MS/MS). Samples were spiked with an internal standard mixture containing phosphatidylcholine (PC 17:0/17:0; Avanti Polar Lipids) and ergosterol (Merck). Lipids were extracted using an acidified methanol–chloroform protocol. Briefly, samples (n = 6 per group) were mixed with HCl, methanol, and chloroform, vortexed, and centrifuged. The organic phase was collected, re-extracted, combined, and evaporated under vacuum. Dried lipids were resuspended in methanol:chloroform (4:1, v/v) and separated on a reverse-phase $C_{18}$ column at 50 °C. The mobile phases consisted of water with 1% formic acid (A) and methanol (B), using a gradient with increasing organic phase. Detection was performed on a triple-quadrupole mass spectrometer (API 2000, Sciex) operated in positive ion mode. Lipid species were quantified using Analyst software (v1.6.2) based on internal standards and external calibration curves.

**Sample preparation for HPLC.** Neuro2a cells were incubated with 10 µM CYM50769 (dissolved in DMEM, high glucose, GlutaMAX supplement, pyruvate growth serum (GIBCO,

#31966021) with 1% DMSO, 10% fetal bovine serum (Biowest, #1810)) for the specified time points. Cells were lysed by freezing at −80 °C. Proteins were subsequently precipitated by addition of ice-cold methanol, followed by centrifugation at 14,000 rpm for 15 min. The supernatant was collected and filtered through a 0.45 µm pore-size filter. Samples were then dried at 40 °C using a vacuum centrifuge. The dried residues were resuspended in 50 µl acetonitrile, incubated in an ultrasonic bath, and briefly vortexed. After a second centrifugation step (14,000 rpm, 15 min), the supernatant was transferred to HPLC vials for analysis.

**Targeted and untargeted metabolomics.** Metabolite profiling of CYM50769 in Neuro2A-cells was performed using ultra-high-performance liquid chromatography coupled to high-resolution mass spectrometry (UHPLC–HRMS). Samples were separated on a $C_{18}$ column (Accucore, 100 × 2.1 mm, 2.6 µm) using a binary solvent system consisting of water with 0.1% formic acid and acetonitrile at a flow rate of 0.4 ml $min^{-1}$. The autosampler was maintained at 10 °C and the column at 25 °C. Mass spectra were acquired on an Orbitrap Exploris 480 mass spectrometer (Thermo Fisher Scientific) equipped with a heated electrospray ionization source. Data were collected in positive and negative ionization modes over a mass range of 100–1500 *m/z* at a resolution of 180,000. To minimize source contamination, LC flow was diverted to waste during the initial and final phases of the run. Relative quantification of CYM50769 was performed by integration of the protonated molecular ion (*m/z* 417.1001 ± 3 ppm) using Xcalibur (v4.5) and FreeStyle (v1.8SP2). Peak detection was based on extracted ion chromatograms using standard parameters.
Liver samples for lipid metabolomics (n = 6 per group) were prepared using a modified protocol from [2]. Briefly, the frozen tissues were weighed and homogenized using a cell homogenizer in ice-cold methanol-water (4 ml methanol per gram of tissue and 0.85 ml water per gram) with 5 silicon carbide fragments (2–3 mm particle size). 4 ml/g of dichloromethane and 2 ml/g of water were added to each homogenate. The samples were vortexed and then placed at -20 °C for 15 min. After extraction, the samples were centrifuged at 4 °C and 12,000 rpm for 15 min. The upper methanol/water phase (containing polar metabolites) was transferred to a 1.5 ml vial, dried using a SpeedVac, and then subjected to LC-HRMS analysis. Liver weight was used as to normalize metabolite abundance for statistical data analysis.

**Untargeted lipid metabolomics data analysis.** 40 curated metabolites were identified by spectral matching using SIRIUS. These correspond to confidence level 2 annotations [3]. Data-dependent MS/MS ($ddMS^2$) acquisition was performed in positive and negative polarity, with 5 dependent scans, isolation window of *m/z* 1.5, energy of collision (20, 60), maximum injection time of 100 ms and orbitrap resolution of 30,000 for each MS/MS spectrum. MS/MS spectra enabled advanced metabolite annotation through spectral similarity matching with databases using SIRIUS implemented within Compound Discoverer 3.5 (Thermo Fisher Scientific). Remaining metabolites were assigned putative annotations (confidence level 3 with partial match, and confidence level 2 for full match). 2,278 features were detected, among which 1,213 were found significantly regulated across treatments. From these, 40 metabolites were annotated at confidence level 2 with full spectral matches and were retained in the curated metabolites list. Raw data have been deposited in the MassIVE database (MSV000101643).

**CVS and behavior.** The CVS paradigm was performed as described previously [4]. Briefly, mice were subjected to 21 consecutive days of stress exposure consisting of three stressors applied in a semi-random sequence. The stressors included 1 h of tube restraint, tail suspension, or exposure to 100 mild, randomly delivered foot shocks.

Behavioral testing was carried out during the animals' active (dark) phase under red light conditions, whereas stress procedures were applied during the first half of the light phase. Animal well-being was monitored throughout, with no immediate adverse effects observed. Body weight was recorded at least weekly. Mice were group-housed (2 to 5 per cage), except for temporary single housing during behavioral testing. The forced swim test was performed as described previously [5,6].

**Statistics and graphics.** Statistical analysis was performed using GraphPad Prism. Two-tailed Student's t-tests were used for comparisons between two groups., with Welch correction applied where appropriate. Nonparametric data were analyzed using the Mann-Whitney test. For comparisons involving one or two factors, one-way or two-way ANOVA followed by Tukey's or Bonferroni *post hoc* tests were used, respectively. For combined analyses across sexes, data were normalized to sex-matched controls. All behavioral experiments were conducted with the experimenter blinded to group allocation.

**Data visualization and overlap analyses.** Heatmaps display significantly altered gene products ordered by $\log_2$ fold change. Corresponding gene products from reference groups, irrespective of their significance, are included for comparison. Overlaps between significant gene sets are visualized using Venn diagrams (circle sizes not to scale). Functional enrichment analyses were performed using Metascape as described previously [7]. To assess concordance in protein regulation across datasets, rank-rank hypergeometric overlap (RRHO) analysis was performed [8,9]. Gene products were ranked based on adjusted P values with directionality defined by fold change. Pairwise comparisons were evaluated using hypergeometric testing, and multiple testing correction was applied using the Benjamini–Yekutieli false discovery rate method.

**Chord diagram analysis.** Chord diagrams were generated using R (v4.4.3) with the packages tidyverse (v2.0.0) [10] and circlize (v0.4.16) [11]. Proteomics data were imported using readxl (v1.4.5) [12] and filtered to include significant DEPs ($q < 0.05$) excluding keratin contaminants. For each comparison, shared proteins between tissue pairs were identified based on UniProt IDs, and the mean $\log_2$ fold change was calculated. Chord diagrams were generated using CairoPDF (v1.6.5) [13], where connections represent shared DEPs between tissues. Chord thickness reflects the number of shared proteins, and color indicates the average direction and magnitude of regulation.

**Table S1: Metrics on DEPs in mice.**

| Organ | Parameter | Female "Stress" | Female "CYM50769" | Male "Stress" | Male "CYM50769" |
|---|---|---|---|---|---|
| Adipose tissue | total sign. (no NAN/Krt) | 825 | 940 | 1213 | 876 |
| | % up | 29.33 | 66.49 | 42.54 | 52.51 |
| | Total detected DEPs | 4730 | 4732 | 4737 | 4747 |
| | % sign. changed | 17.44 | 19.86 | 25.61 | 18.45 |
| Adrenal gland | total sign. (no NAN/Krt) | 1438 | 1368 | 902 | 768 |
| | % up | 37.76 | 73.54 | 30.16 | 72.66 |
| | Total detected DEPs | 6334 | 6325 | 6220 | 6222 |
| | % sign. changed | 22.70 | 21.63 | 14.50 | 12.34 |
| Heart | total sign. (no NAN/Krt) | 321 | 549 | 464 | 563 |
| | % up | 36.76 | 62.11 | 58.19 | 57.02 |
| | Total detected DEPs | 3370 | 3361 | 3372 | 3363 |
| | % sign. changed | 9.53 | 16.33 | 13.76 | 16.74 |
| Hypothalamus | total sign. (no NAN/Krt) | 255 | 101 | 306 | 217 |
| | % up | 82.35 | 43.56 | 39.54 | 31.34 |
| | Total detected DEPs | 6183 | 6196 | 6228 | 6227 |
| | % sign. changed | 4.12 | 1.63 | 4.91 | 3.48 |
| Kidney | total sign. (no NAN/Krt) | 350 | 497 | 264 | 176 |
| | % up | 62.29 | 61.17 | 69.70 | 40.34 |
| | Total detected DEPs | 5983 | 5983 | 6156 | 6156 |
| | % sign. changed | 5.85 | 8.31 | 4.29 | 2.86 |
| Liver | total sign. (no NAN/Krt) | 781 | 227 | 302 | 423 |
| | % up | 46.48 | 52.42 | 54.64 | 59.10 |
| | Total detected DEPs | 4936 | 4945 | 5334 | 5326 |
| | % sign. changed | 15.82 | 4.59 | 5.66 | 7.94 |
| Lung | total sign. (no NAN/Krt) | 454 | 474 | 938 | 514 |
| | % up | 32.60 | 76.79 | 80.06 | 18.09 |
| | Total detected DEPs | 6135 | 6133 | 6128 | 6116 |
| | % sign. changed | 7.40 | 7.73 | 15.31 | 8.40 |
| NAc | total sign. (no NAN/Krt) | 172 | 55 | 312 | 205 |
| | % up | 74.42 | 45.45 | 45.51 | 51.22 |
| | Total detected DEPs | 6381 | 6381 | 6345 | 6348 |
| | % sign. changed | 2.70 | 0.86 | 4.92 | 3.23 |
| Gonads | total sign. (no NAN/Krt) | 996 | 1303 | 362 | 431 |
| | % up | 55.02 | 56.02 | 67.40 | 59.16 |
| | Total detected DEPs | 6730 | 6757 | 7448 | 7461 |
| | % sign. changed | 14.80 | 19.28 | 4.86 | 5.78 |
| PFC | total sign. (no NAN/Krt) | 125 | 387 | 46 | 88 |
| | % up | 56.00 | 40.05 | 43.48 | 25.00 |
| | Total detected DEPs | 6215 | 6210 | 6273 | 6278 |
| | % sign. changed | 2.01 | 6.23 | 0.73 | 1.40 |
| Pituitary gland | total sign. (no NAN/Krt) | 780 | 1766 | 277 | 291 |
| | % up | 35.64 | 82.96 | 20.58 | 49.48 |
| | Total detected DEPs | 7554 | 7574 | 6893 | 6890 |
| | % sign. changed | 10.33 | 23.32 | 4.02 | 4.22 |

| | | | | | |
|---|---|---|---|---|---|
| Small Intestine | total sign. (no NAN/Krt) | 721 | 1237 | 1827 | 862 |
| | % up | 68.10 | 56.83 | 30.87 | 66.36 |
| | Total detected DEPs | 6337 | 6330 | 6466 | 6450 |
| | % sign. changed | 11.38 | 19.54 | 28.26 | 13.36 |
| Serum | total sign. (no NAN/Krt) | 224 | 151 | 167 | 142 |
| | % up | 30.36 | 84.11 | 77.84 | 28.87 |
| | Total detected DEPs | 679 | 671 | 672 | 663 |
| | % sign. changed | 32.99 | 22.50 | 24.85 | 21.42 |

**Table S2: Metrics on human serum samples**

| | Significance | CTRL Women [1] | CTRL Men [1] | MDD Women [1] | MDD Men [1] |
|---|---|---|---|---|---|
| Group size | | 30 | 40 | 33 | 49 |
| Age | | 35.83 ± 10.38 | 33.80 ± 9.78 | 36.79 ± 12.21 | 33.04 ± 9.70 |
| BMI | | 24.38 ± 4.38 | 24.23 ± 2.92 | 24.31 ± 5.69 | 24.26 ± 3.64 |
| Waist-to-hip-ratio | ***effect of MDD and sex [2] | 0.86 ± 0.05 | 0.91 ± 0.07 | 0.90 ± 0.05 | 0.94 ± 0.06 |
| Cigarettes / day | ***effect of MDD, *effect of sex, *interaction [2] | 2.53 ± 4.89 | 2.13 ± 5.18 | 4.52 ± 8.64 | 10.88 ± 10.27 |
| Symptom onset before admission (months) | | | | 35.30 ± 67.74 | 68.58 ± 89.00 |
| GAF score | | | | 49.19 ± 11.57 | 48.27 ±12.75 |
| HAMD score | | | | 21.39 ± 5.93 | 20.20 ± 6.10 |
| HAMD1 (depr. mood) | P = 0.0355* [3] | | | 3.15 ± 0.62 | 2.78 ± 0.87 |
| HAMD2 (feelings of guilt) | | | | 1.61 ± 0.75 | 1.49 ± 0.84 |
| HAMD3 (suicide) | | | | 1.30 ± 1.40 | 1.78 ± 1.43 |
| HAMD4 (diffic. falling asleep) | | | | 1.06 ± 0.90 | 1.29 ± 0.91 |
| HAMD5 (diffic. staying asleep) | | | | 1.09 ± 0.88 | 0.90 ± 0.92 |
| HAMD6 (morning sleep distu.) | | | | 1.00 ± 0.94 | 0.86 ± 0.98 |
| HAMD7 (work, other activities) | | | | 2.45 ± 1.23 | 2.78 ± 1.16 |
| HAMD8 (depr. inhibition) | | | | 0.21 ± 0.65 | 0.12 ± 0.33 |
| HAMD9 (agitation) | | | | 0.45 ± 0.62 | 0.55 ± 0.71 |
| HAMD10 (anxiety psycholog.) | | | | 2.76 ± 1.56 | 2.04 ± 1.74 |
| HAMD11 (anxiety somatic) | | | | 0.91 ± 1.26 | 0.67 ± 1.09 |
| HAMD12 (physical symptoms, gastrointestinal) | | | | 0.33 ± 0.54 | 0.35 ± 0.63 |
| HAMD13 (physical symptoms, general) | | | | 0.76 ± 0.90 | 0.69 ± 0.82 |
| HAMD14 (genital symptoms) | | | | 0.67 ± 0.85 | 0.55 ± 0.82 |
| HAMD15 (hypochondria) | | | | 0.30 ± 0.77 | 0.18 ± 0.60 |
| HAMD16a (weight loss from history) | | | | 0.64 ± 0.86 | 0.69 ± 0.92 |
| HAMD16b (weight loss assessed in clinic) | | | | 0.42 ± 0.71 | 0.59 ± 0.84 |
| HAMD17 (insight into illness) | | | | 0.03 ± 0.17 | 0.00 ± 0.00 |
| HAMD18a (daily fluctuations morning / evening) | | | | 0.73 ± 0.80 | 0.69 ±0.80 |
| HAMD18b (daily fluctuations intensity) | | | | 0.94 ± 0.97 | 0.88 ± 0.95 |
| HAMD19 (depersonalization, | | | | 0.12 ± 0.42 | 0.08 ± 0.34 |

| | | | | | |
|---|---|---|---|---|---|
| derealization) | | | | | |
| HAMD20 (paranoid sympt.) | P = 0.0304* [3] | | | 0.36 ± 0.60 | 0.12 ± 0.39 |
| HAMD21 (OC symptoms) | | | | 0.09 ± 0.29 | 0.12 ± 0.39 |
| BDI | | | | 30.97 ± 9.84 | 30.37 ± 8.85 |
| BdiI01 (sadness) | | | | 1.56 ± 0.76 | 1.45 ± 0.82 |
| BdiI02 (pessimism) | | | | 1.44 ± 0.88 | 1.47 ± 1.00 |
| BdiI03 (past failures) | | | | 1.28 ± 1.05 | 1.55 ± 0.94 |
| BdiI04 (loss of pleasure) | | | | 2.06 ± 0.80 | 2.08 ± 0.73 |
| BdiI05 (feelings of guilt) | | | | 1.38 ± 0.98 | 1.22 ± 1.03 |
| BdiI06 (feelings of being punished) | | | | 0.97 ± 0.93 | 0.67 ± 0.85 |
| BdiI07 (self-dislike) | | | | 1.47 ± 0.88 | 1.71 ± 1.02 |
| BdiI08 (self-blame) | | | | 1.56 ± 0.98 | 1.63 ± 0.81 |
| BdiI09 (suicidal thoughts or wishes) | P = 0.0187* [3] | | | 0.78 ± 0.55 | 1.20 ± 0.89 |
| BdiI10 (crying) | | | | 1.41 ± 0.87 | 1.63 ± 1.18 |
| BdiI11 (restlessness) | | | | 1.34 ± 0.87 | 1.29 ± 0.98 |
| BdiI12 (lack of interest) | | | | 1.53 ± 0.80 | 1.76 ± 0.95 |
| BdiI13 (indecisiveness) | | | | 1.91 ± 0.96 | 1.51 ± 1.04 |
| BdiI14 (worthlessness) | | | | 1.66 ± 0.65 | 1.45 ± 0.96 |
| BdiI15 (loss of energy) | | | | 1.81 ± 0.74 | 1.73 ± 0.78 |
| BdiI16 (changes in sleep patterns) | P = 0.0220* [3] | | | 2.09 ± 0.64 | 1.67 ± 0.88 |
| BdiI17 (irritability) | | | | 1.22 ± 0.97 | 0.98 ± 0.97 |
| BdiI18 (changes in appetite) | | | | 1.44 ± 1.05 | 1.35 ± 0.95 |
| BdiI19 (difficulty concentrating) | | | | 1.47 ± 0.76 | 1.61 ± 0.73 |
| BdiI20 (fatigue) | | | | 1.34 ± 0.97 | 1.41 ± 1.02 |
| BdiI21(loss of interest in sex) | | | | 1.25 ± 1.11 | 0.98 ± 0.99 |

*Legend:* [1] Sex information was based on self-identification. [2] Two-way ANOVA & Bonferroni *post hoc* test; wait-to-hip ratio: effect of MDD: $F_{(1,148)}$ = 12.88, ***P = 0.0005; effect of sex: $F_{(1,148)}$ = 21.29, ***P ( 0.0001. *Post hoc* test: Effect of MDD in women: P < 0.05, men: P < 0.05; effect of sex in controls: P < 0.01, in MDD: P < 0.01. Cigarettes per day (self-reported): effect of MDD: $F_{(1,148)}$ = 17.03, ***P < 0.0001; effect of sex: $F_{(1,148)}$ = 5.25, *P = 0.0234; interaction MDD and sex: $F_{(1,148)}$ = 6.75, *P = 0.0103. *Post hoc* test: Effect of MDD in men: P < 0.001; effect of sex in MDD: P < 0.001. [3] Student's t-test (2-tailed); only significant data are reported. GAF – global assessment of function; HAMD – Hamilton depression rating scale; depr. – depressive, diffic. – difficulty, distu. – disturbances, psycholog. – psychological, sympt. – symptoms,OC – obsessive compulsive; BDI – Beck depression inventory; BdiI - Beck depression inventory, second edition

**Table S3. Gradient for UHPLC / HRMS measurement.**

| time [min] | solvent B [%] |
|---|---|
| 0 | 0 |
| 0.3 | 0 |
| 7.7 | 100 |
| 11.0 | 100 |
| 11.1 | 0 |
| 13.0 | 0 |

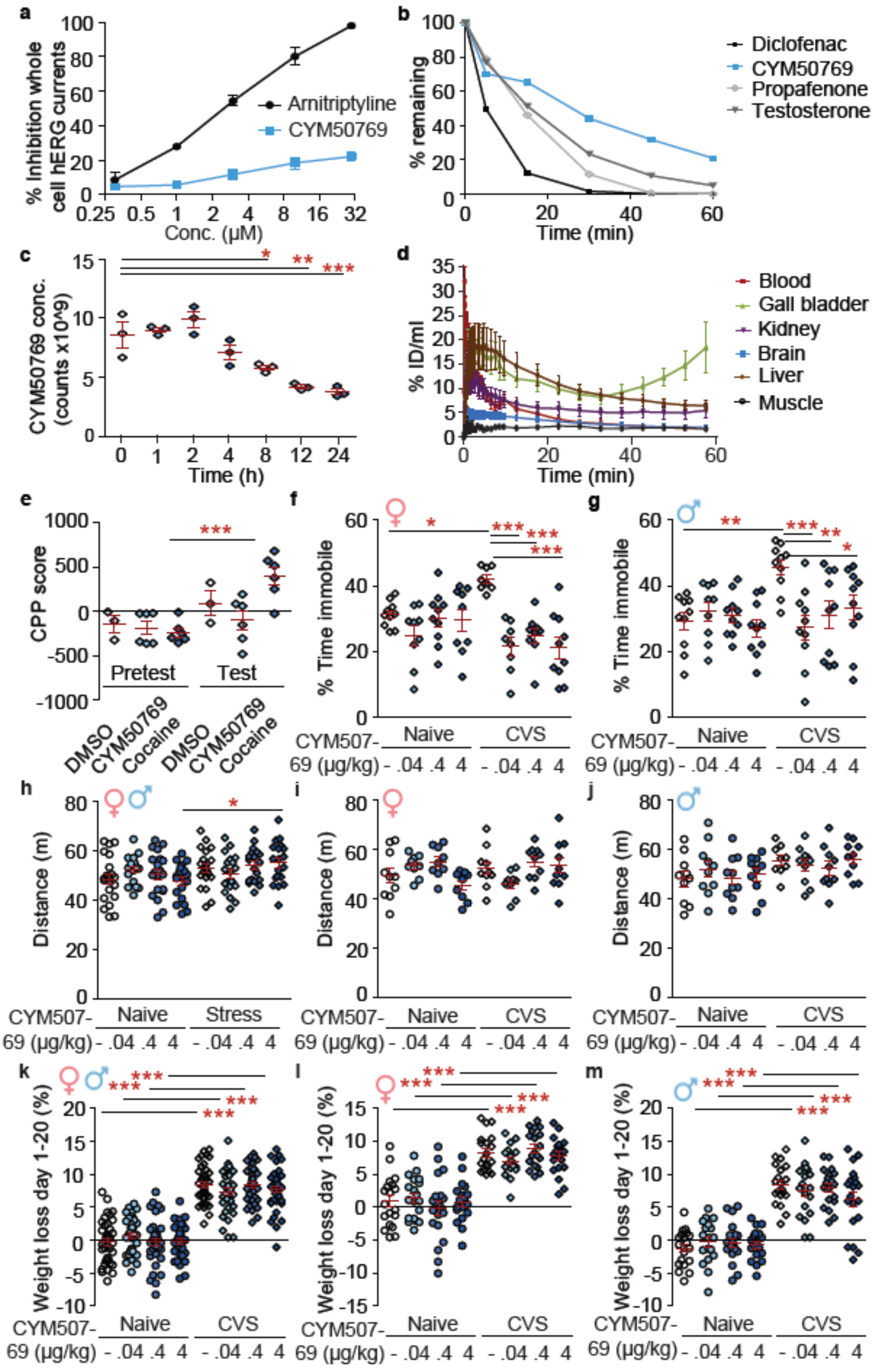
a
% Inhibition whole cell hERG currents
Arnitriptyline
CYM50769
Conc. (μM)
b
% remaining
Time (min)
Diclofenac
CYM50769
Propafenone
Testosterone
c
CYM50769 conc. (counts x10^9)
Time (h)
d
% ID/ml
Time (min)
Blood
Gall bladder
Kidney
Brain
Liver
Muscle
e
CPP score
Pretest
Test
DMSO
CYM50769
Cocaine
f
% Time immobile
Naive
CVS
CYM507-69 (μg/kg)
- .04 .4 4
g
% Time immobile
h
Distance (m)
Naive
Stress
i
Distance (m)
j
Distance (m)
k
Weight loss day 1-20 (%)
l
Weight loss day 1-20 (%)
m
Weight loss day 1-20 (%)

**Fig. S1 | Medicinal chemistry and animal safety. a**, hERG channel assays. **b**, Liver microsomal stability assays. **c**, HPLC shows stability of CYM50769 for at least 8h in cell culture. Neuro2A-cells were incubated in growth medium containing CYM50769 at 37 °C. At certain time points, cells and medium were collected. And CYM50769 levels determined. n = 3 per group. One-way ANOVA: F = 18.4, P < 0.0001. Tukey *post hoc test*: Only comparison to 0 h time point shown. 0 h vs 8 h: *P < 0.05; 0 h vs 12 h: **P < 0.01; 0 h vs 24 h: ***P < 0.001. **d**, PET imaging: Distribution of CYM50769 across organs and time. IG: Injected Dose per ml. **e**-**m**, Statistics: 2-way ANOVA & Bonferroni *post hoc* test. **e**, Individual CPP scores during pre-test and test show a cocaine-induced place preference but no conditioning for CYM50769. n = 3, 6, 6 per group. interaction test by drug: $F_{(2,24)}$ = 4.00; *post hoc* test: pretest vs. test within cocaine group: *** P < 0.001; other groups: n.s. **f**, **g**, Forced swim test data separated by sex. In both sexes, CVS increases the immobility time, and all doses of CYM50769 rescue this effect. **f**, Females: n = 10, 9, 10, 9, 8, 8, 9, 10; interaction stress by CYM50769 concentration: $F_{(3,65)}$ = 5.35; P < 0.001; effect of CYM50769 concentration: $F_{(3,65)}$ = 10.38; P < 0.0001. *Post hoc* test: stress effect in vehicle group: *P < 0.05; effect of low, moderate and high CYM50769 doses versus vehicle group within CVS-treated mice: all ***P < 0.001. **g**, Males: n = 10, 9, 10, 10, 10, 10, 10, 10; interaction stress by CYM50769 concentration: $F_{(3,71)}$ = 4.36; P < 0.001; effect of CVS: $F_{(1,71)}$ = 4.19; P < 0.05. *Post hoc* test: stress effect in vehicle group: **P < 0.01; effect of low CYM50769 dose versus vehicle group within CVS-treated mice: ***P < 0.01; moderate CYM50769 vs. vehicle within CVS-group**:** **P <0.01; high CYM50769 vs. vehicle within CVS group: P < 0.05. **h-j**, Distance travelled in an open field arena shows no altered locomotor activity at moderate CYM50769 doses. **h,** Both sexes combined. n = 20, 17, 20, 19, 20, 20, 19, 19; effect of CVS: $F_{(1,146)}$ = 7.1; P = 0.0087; *post hoc* test: CVS effect within highest dose of CYM50769: P < 0.05. **i**, Females. n= 10, 9, 9, 9, 10, 9, 10, 10; interaction between CVS and CYM50769: $F_{(1,68)}$ = 3.6; P = 0.0182; *post hoc* test: n.s. **j**, Males. n = 10, 10,10, 10, 9, 10, 10, 10; effect of CVS: $F_{(1,71)}$ = 6.3; P = 0.0143; *post hoc* test: n.s. **k**-**m**, CVS, but not CYM50769, affects the weight of mice. **k**, Both sexes combined. n = 38, 35, 38, 35, 39, 36, 38, 37; effect of CVS: $F_{(1,288)}$ = 498.3; P < 0.001. *Post hoc* test: stress effect in all treatment groups: ***P < 0.001. **l**, Females. n = 20, 19, 20, 19, 20, 18, 20, 20; effect of CVS: $F_{(1,148)}$ = 149.0; P < 0.0001. *Post hoc* test: stress effect in all treatment groups: ***P < 0.001. **m**, Males. n = 19, 18, 20, 18, 20, 19, 19, 20; effect of CVS: $F_{(1,145)}$ = 223.7; P < 0.0001. *Post hoc* test: stress effect in all treatment groups: ***P < 0.001.

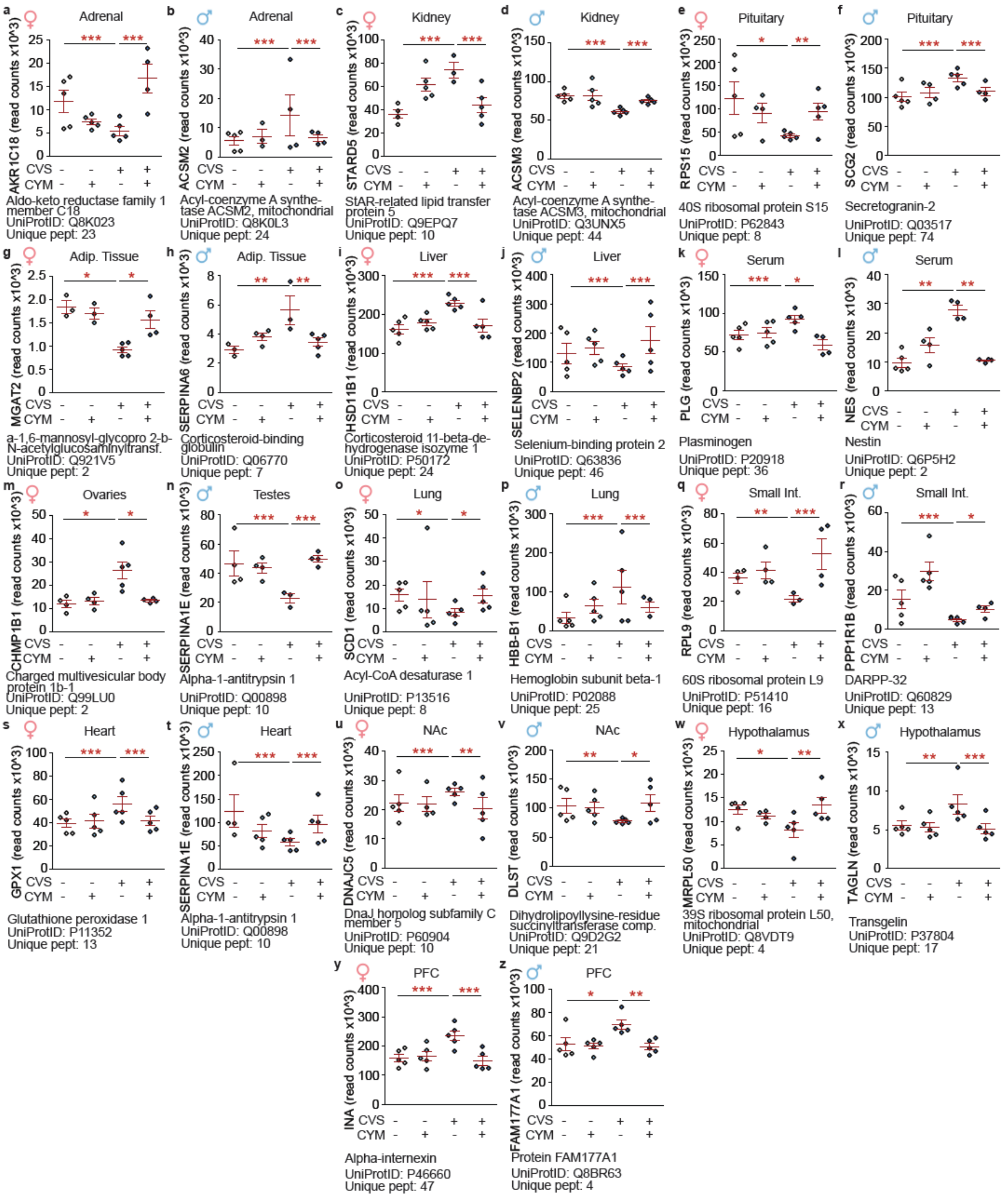


**Fig. S2 | Examples of DEPs that were affected by stress and NPBWR1 inhibition.** Proteomics raw data are shown. Statistics refer to adjusted P-value from Spectronaut (*P < 0.05; ** P < 0.01, ***P < 0.001). CYM – CYM50769.

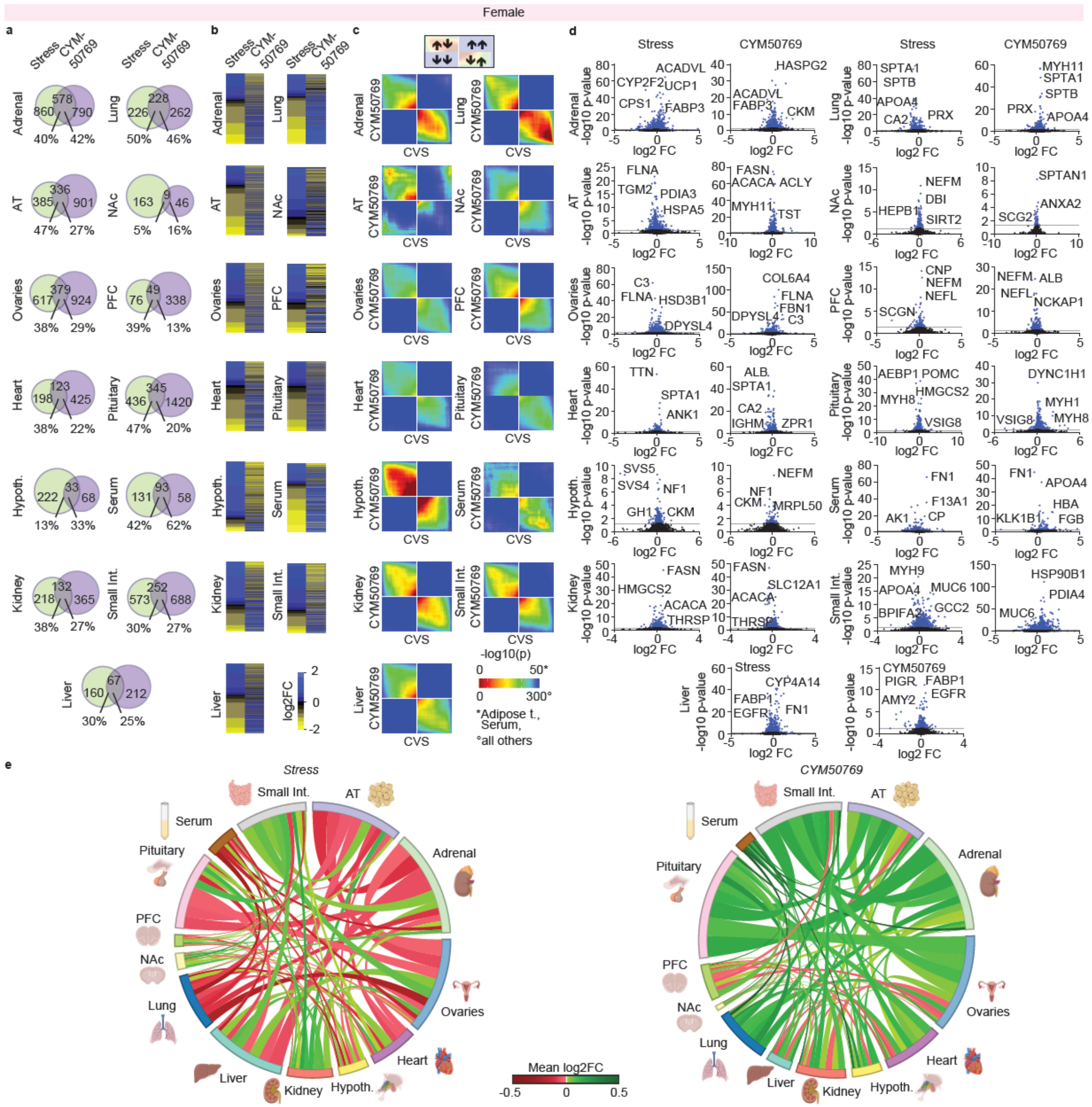


**Fig. S3 | CYM50769 reverses systemic effects of chronic stress in female mice. a**, Overlap between DEPs significantly changed by stress and CYM50769 (Venn diagrams), shows partial reversal of stress effects by CYM50769. **b**, Heatmaps depict DEPs that are significantly altered by stress and were sorted by $\log_2$ fold change within the stress group, with the same DEPs depicted for CYM50769 in the right lanes regardless of the adjusted P-value in that group. The heatmaps show an inverse $\log_2$ fold change within the CYM50769-group across sexes and tissues. **c**, RRHO analysis shows inverse correlation between CVS and CYM50769 across sexes and tissues. **d**, Volcano plots show the distribution of significance and $\log_2$ fold change altered DEPs. **e**, Chord diagrams reveal stress-induced changes in organ connectomics. These are, at least in part, reversed by CYM50769. “Stress”: DMSO Naïve vs. DMSO CVS; “CYM50769”: DMSO CVS vs. CYM50769 CVS. Sketches of organs were made with biorender.com.

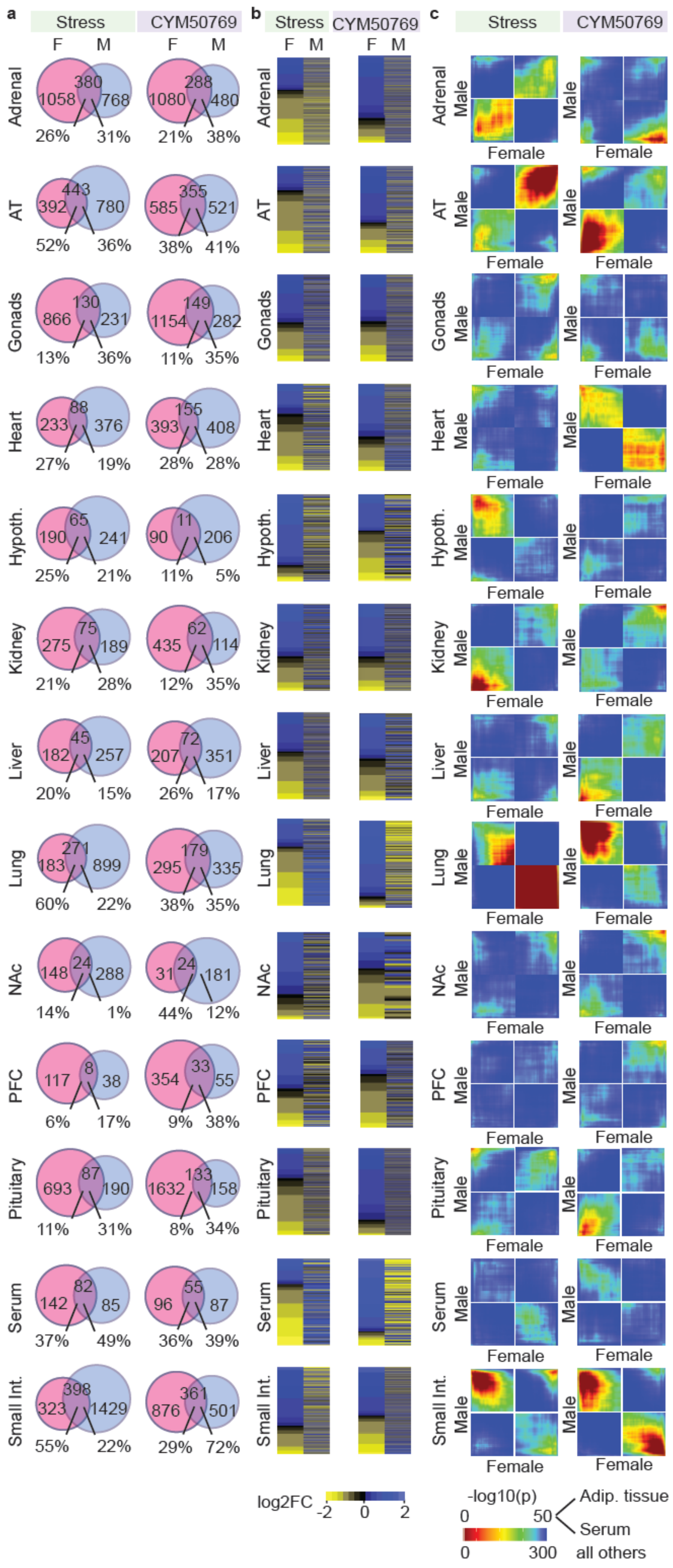
a
Stress
CYM50769
F M
Adrenal
1058 380 768
26% 31%
1080 288 480
21% 38%
AT
392 443 780
52% 36%
585 355 521
38% 41%
Gonads
866 130 231
13% 36%
1154 149 282
11% 35%
Heart
233 88 376
27% 19%
393 155 408
28% 28%
Hypoth.
190 65 241
25% 21%
90 11 206
11% 5%
Kidney
275 75 189
21% 28%
435 62 114
12% 35%
Liver
182 45 257
20% 15%
207 72 351
26% 17%
Lung
183 271 899
60% 22%
295 179 335
38% 35%
NAc
148 24 288
14% 1%
31 24 181
44% 12%
PFC
117 8 38
6% 17%
354 33 55
9% 38%
Pituitary
693 87 190
11% 31%
1632 133 158
8% 34%
Serum
142 82 85
37% 49%
96 55 87
36% 39%
Small Int.
323 398 1429
55% 22%
876 361 501
29% 72%
b
log2FC -2 0 2
c
Male
Female
-log10(p)
0 50
Adip. tissue
Serum
0 300
all others

**Fig. S4 | Sex-specific proteome changes in response to chronic stress and CYM50769 treatment. a**, Percentage of significantly regulated DEPs in tissues of male versus female mice following stress and CYM50769. Venn diagrams show moderate but comparable overlap of stress- and CYM50769-regulated DEPs across sexes. Sex-specificity in stress response varies across tissues with the smallest overlap in the brain regions NAc, PFC and pituitary gland (each < 20%). **b**, Heatmaps depict the directionality of stress- and CYM50769-induced changes in tissue proteomes across sexes. Most tissues show no common directionality of stress-induced changes across sexes, with the hypothalamus, lung, and serum demonstrating a reverse association. **c**, RRHO plots show a moderate correlation between stress-effects across sexes in the adrenal gland and adipose tissue, while they reveal an inverse correlation in the hypothalamus, lung, and small intestine.

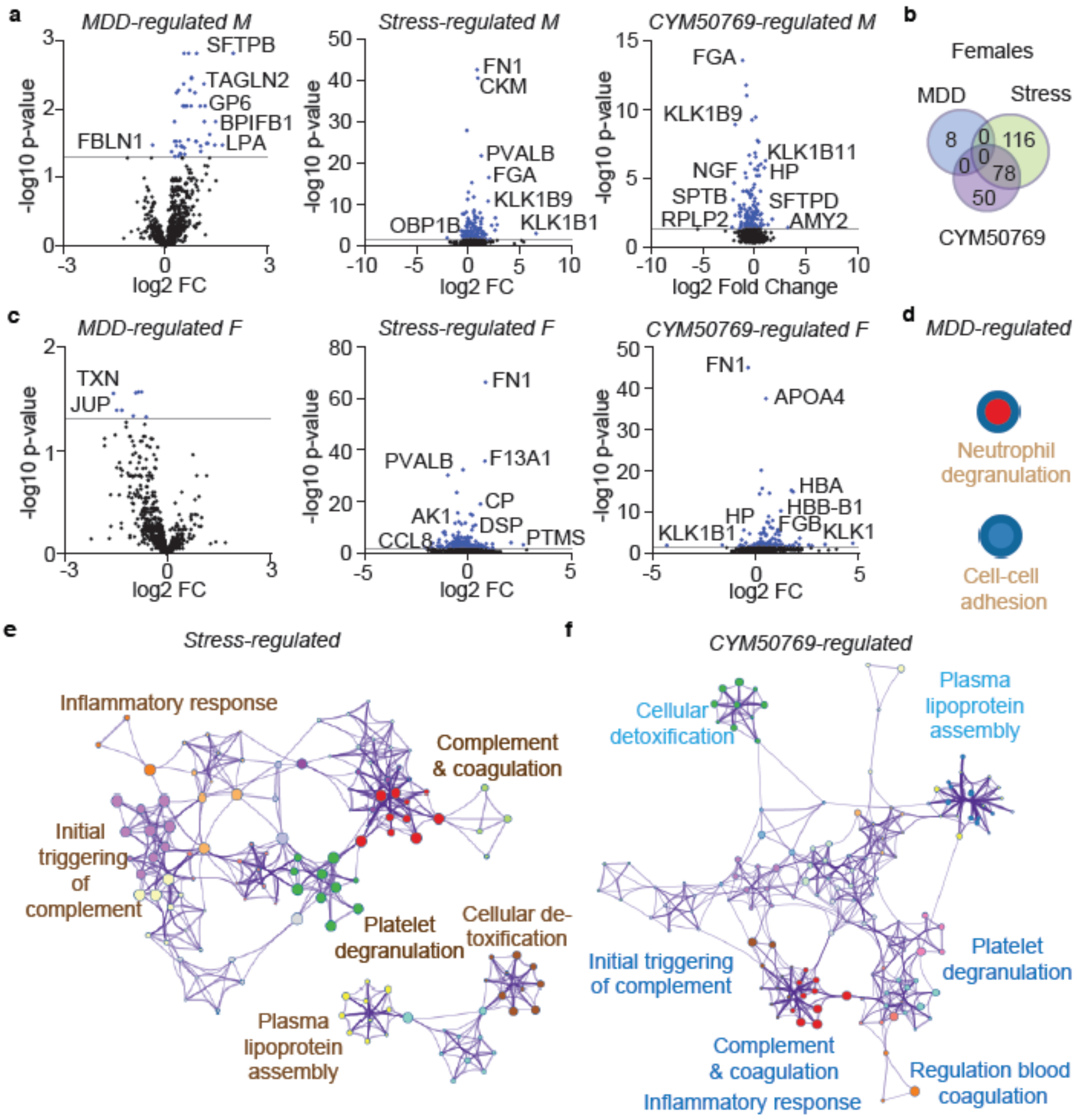


**Fig. S5 l Extended comparisons of serum data from MDD patients and mice receiving CVS or CYM507569 treatment. a**, Volcano plots from male cohorts. From left to right: effects of MDD, CVS, and CYM50769. **b**, Overlapping DEPs from female cohorts that are significantly regulated by MDD, CVS, and CYM50769. **c**, Volcano plots from female cohorts. From left to right: effects of MDD, CVS, and CYM50769. **d-f**, Metascape analyses from serum of female cohorts. Blue: upregulation, brown: downregulation. **d**, Few pathways are significantly altered by MDD. Note that those are downregulated. **e**, Pathways that are downregulated by stress include coagulation, platelet degranulation, and inflammation as well as plasma lipoprotein assembly. **f**, CYM50769-regulated pathways are predominantly upregulated and include coagulation, inflammation, platelet degranulation, and plasma lipoprotein assembly. “Stress” comparison: DMSO Naive vs. DMSO CVS; “CYM50769” comparison: CYM50769 CVS vs. DMSO CVS.

**a** Stress

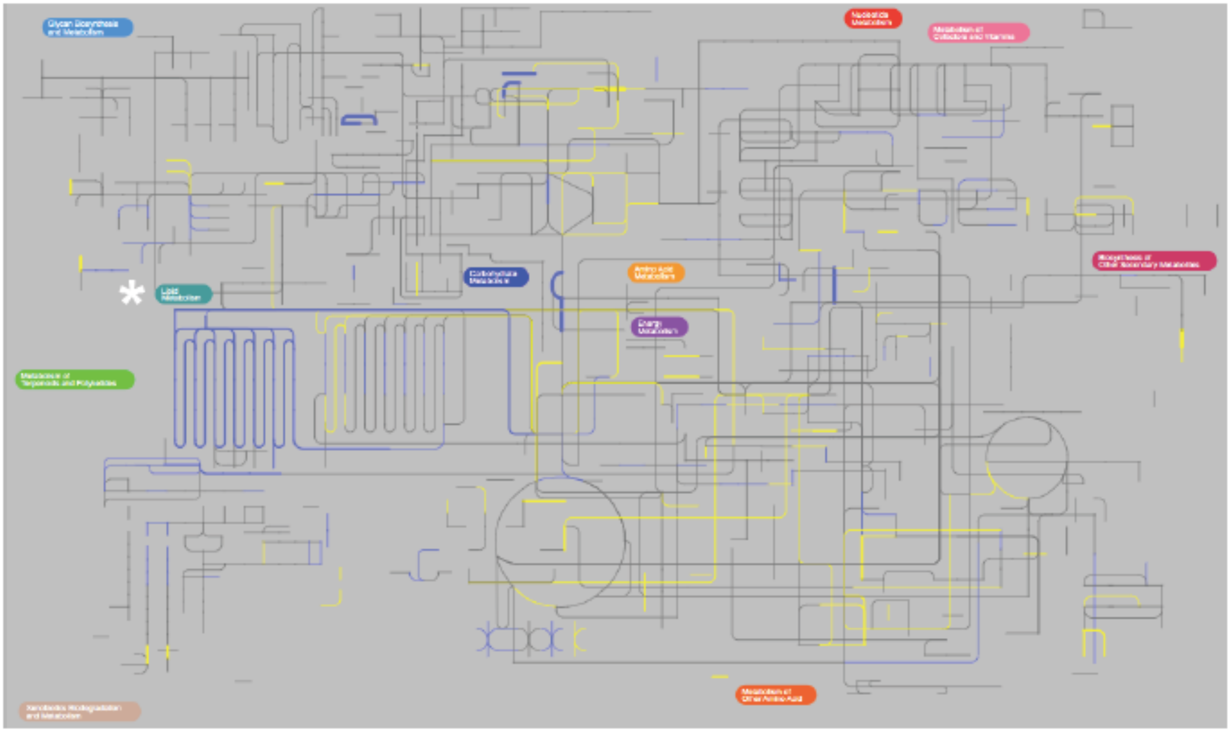

**b** CYM50769

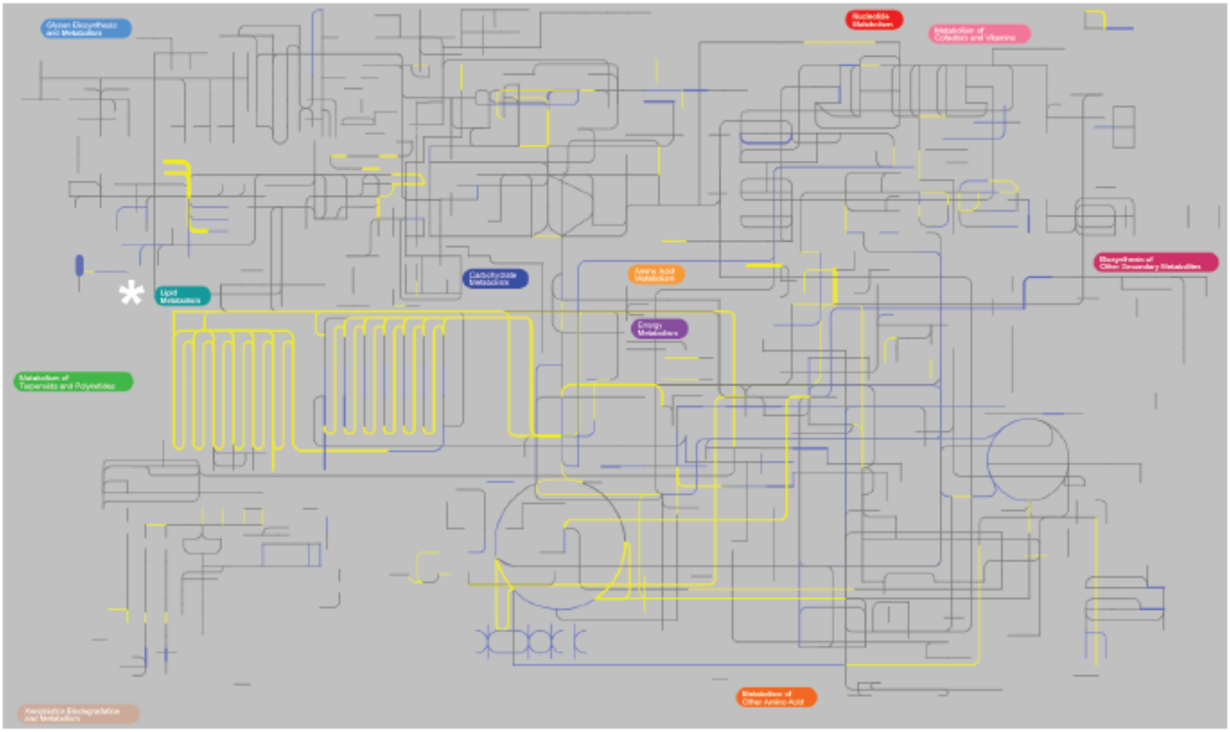

**Fig. S6 | iPath visualization of proteomics data from male liver. a**, Stress effects. **b**, Effects of NPBWR1 inhibition. Asterisk: Lipid metabolism. Blue: upregulation. Yellow: downregulation. iPath visualization for all tissues and sexes can be found on *https://stress-atlas.com*.

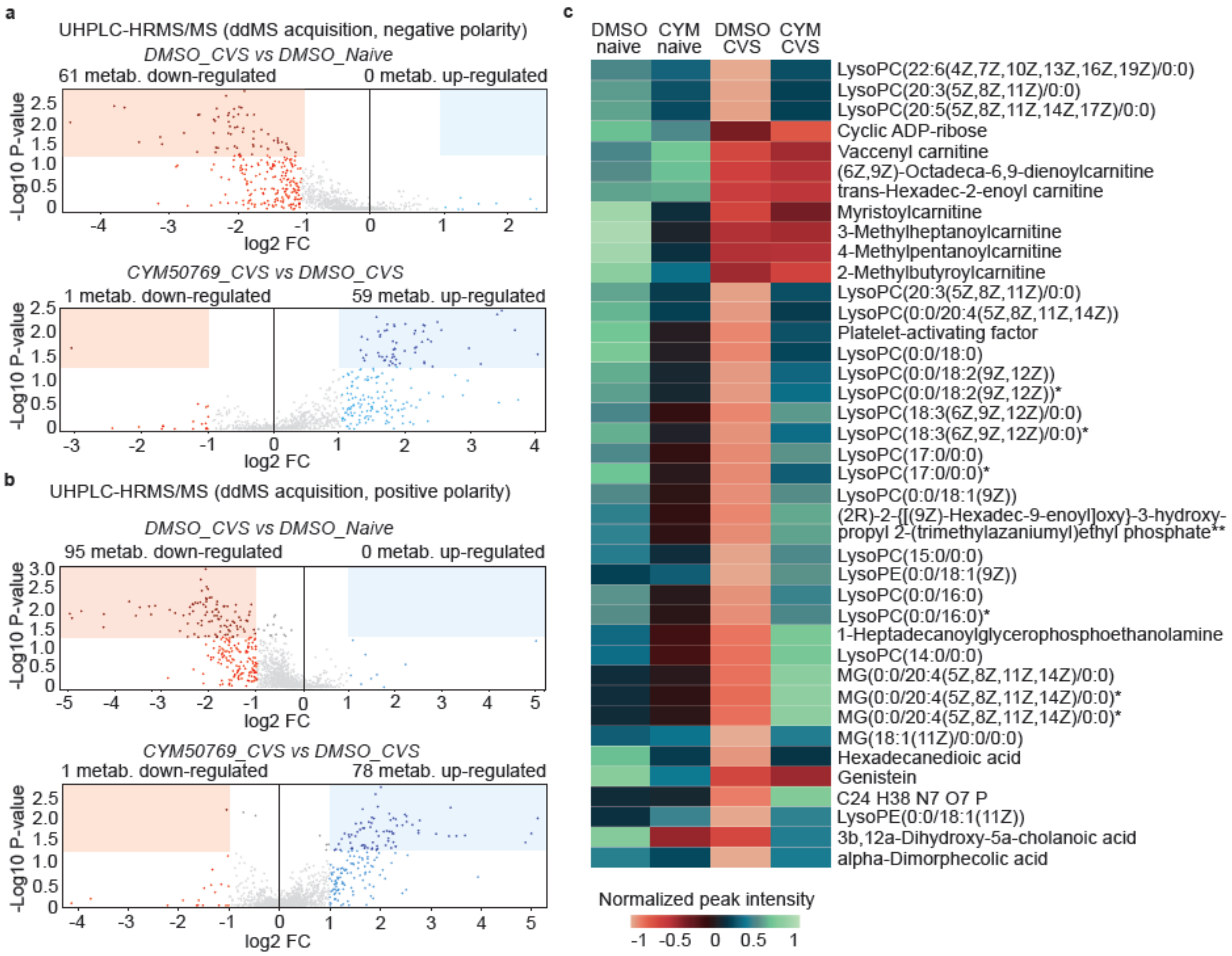


**Fig. S7 | Metabolomics reveal a stress-dependent upregulation of lipid species with partial reversal through NPBWR1 inhibition. a**, **b**, Detected metabolites. These include level 1 assignment (including metabolites that did not match a database). **a**, Metabolites wth negative polarity. **b**, Metabolites with negative polarity. **c**, Heatmap from **Fig. 5** with metabolite names. Note that the blue cluster in lane 4 relates to carnitine signaling, while the majoity of reversed metabolites in lane 4 relates to phosphatidyl cholines (PC) and monoacylglycerols (MG). * isomer of above mentioned metabolite. ** second isomer name was omitted.
